# The evolution of p53 network behavior

**DOI:** 10.1101/098376

**Authors:** Hari Sivakumar, João P. Hespanha, Kyoungmin Roh, Stephen R. Proulx

**Author notes:** All authors contributed extensively to the preparation of this manuscript. A.K.R. and S.R.P per-formed the evolutionary analysis. H.S. and J.PH. developed the algorithm and performed the mathematical analysis. The authors have no conflicts of interest to declare.

## Abstract

We study the evolution of the *p*53 core regulation network across the taxonomic span of humans to protozoans and nematodes. We introduce a new model for the core regulation network in mammalian cells, and conduct a formal analysis of the different network configurations that emerge in the evolutionary path to complexity. Solving the high dimensional equations associated with this model is typically challenging, and we develop a novel algorithm to overcome this problem. A key technical tool used is the representation of the distinct pathways in the core regulation networks as “modules”, such that the behavior of the composite of two or more modules can be inferred from the characteristics of each of the individual modules. Apart from simplifying the complexity of the algorithm, this modular representation also allows us to qualitatively compare the distinct types of switching behaviors each network can exhibit. This then allows us to demonstrate how our model for the core regulation network in mammalian cells matches experimentally observed phenomena, and contrast this with the plausible behaviors admitted by the network configurations in putative primordial organisms. We show that the complexity of the *p*53 core regulation network in vertebrates permits a range of behaviors that can bring about distinct cell fate decisions not possible in the putative primordial organisms.

**Significance Statement:** The *p*53 protein has been protecting organisms from tumors for a billion years. We study the link between the evolution of the *p*53 network structure and its corresponding tumor suppression strategies. We compare the dynamical behaviors in putative primordial organisms with simple networks with the vertebrate network that contains multiple feedback loops. We show that the vertebrate network, but not the ancestral network, can both repair moderate damage and induce apoptosis if too much damage accumulates, balancing the risk of cancer with the cost of too much cell death. Moreover, the complexity of the vertebrate network allows for adaptation, for example to increase *p*53 network sensitivity, which is consistent with recent research on large mammals.

---

A major cause for the emergence of tumors is genetic mutations that occur when the cellular DNA is damaged and not repaired to its original state. When the mutations interfere with the normal regulation of cell division, then cell proliferation could cause the tumor to become cancerous [1]. While there is abundant evidence that exposure to chemical carcinogens and radiation has had an impact on cancer rates [2], there is also evidence that cancer is a general feature of multi-cellular organisms that has presented a long-standing evolutionary challenge [3].

A key component of the DNA damage response mechanism is the *TP*53 gene which expresses the *p*53 tumor suppressor protein, the so-called “guardian of the genome” [4]. *p*53 and its ancestors have been protecting metazoans from mutations arising from DNA damage for over one billion years [5]. Recent research has shown that *p*53 plays this role by first sensing DNA damage, and then mediating various down-stream processes in response. *p*53 can promote cell survival by upregulating genes that bring about cell-cycle arrest or DNA repair. However, *p*53 can also promote cell inactivation or death by bringing about permanent cell-cycle arrest or cell death [6–9]. How *p*53 levels and dynamics determine different cell fates in response to DNA damage is a crucial component of an organism’s tumor suppression mechanism [10–13]. It is therefore no surprise that a mutation in *p*53 is implicated in over 50% of human cancers [14].

One of the oldest known mechanisms cells employ in response to DNA damage is programmed cell death, also known as apoptosis [15]. Cells with DNA that is damaged above some “apoptotic threshold” level generally display sustained high levels of *p*53 [13, 16, 17], inducing the expression of pro-elimination genes [12, 16–18] that bring about apoptosis, thus limiting the risk that a cell lineage will become cancerous. The underlying behavior can be understood in the context of Statistical Decision Theory by imagining that cells attempt to decide on the truthfulness of the statistical hypothesis: “The damaged DNA can be repaired to its original form”.

It is clear that setting a low apoptotic threshold minimizes the risk of cancer since a cell that experiences potentially cancer-causing DNA damage would go to apoptosis. A feature of such networks is that they abet Type 1 errors. Such errors happen when the statistical null hypothesis is true (i.e., DNA can be repaired to its original form), but the cell is killed anyway. While not ideal, Type 1 errors would still be preferable to Type 2 errors, where a cell carrying a cancer-causing mutation could be allowed to live. A single Type 2 error could lead to lethal cancer, and this is in fact known to be one of the “hallmarks” of cancerous clones [2, 19]. That said, overly responsive networks with a low apoptotic threshold may result in developmental defects, reduced tissue growth, or high metabolic costs to replace the dead cells. A major task for *p*53 and the DNA damage response mechanism is therefore to effectively suppress the onset of tumors by selecting different cell fates that would increase whole organism fitness. This is achieved by trading-off between minimizing the number of Type 1 errors that take place, while also trying to suppress all Type 2 errors.

While *p*53 is a vital component of the damage response network, it acts as part of a network of interacting genes that determine cell fate. Extensive genetic studies have uncovered a complex network of hundreds of genes [20] that cooperate to carry out tumor suppression as soon as damage is registered in the genome [21]. There are a multitude of upstream sensors that detect DNA damage, the transducers that relay the information to *p*53 and the down-stream actuators that are regulated by *p*53 to bring about different cell fates [22]. In addition, the dynamics of *p*53 is known to be governed by its interaction with a set of core regulation network proteins including *MDM*2, *PTEN* and *ARF* through multiple feedback pathways [23]. Mutations in some of the respective genes have even been implicated in human cancers, although not to the extent of *p*53 [24].

It is therefore interesting that while homologs of *TP*53 are inferred to have been present in the earliest metazoan ancestors [5], the same cannot necessarily be said about *MDM*2, which is another core regulation protein in the mammalian *p*53 pathway. Based on these results, we introduce the underlying network configurations that are present in extant vertebrates and are inferred to have been present in the ancestors of these vertebrates. We also introduce a novel dynamical model that mathematically describes these networks. Our focus is on whether these alternative network structures would allow for qualitatively distinct forms of *p*53 regulation, which in turn would lead to qualitatively different strategies for dealing with the risk of cancer. Clearly, an individual’s fitness will be low if it is at a high risk of developing fatal cancer. However, a policy whereby *p*53 and its core regulation proteins bring about apoptosis of too many cells at the slightest sign of DNA damage would also incur high metabolic costs.

A core question in the evolution of tumor suppression mechanisms therefore, is how the relevant genetic networks can optimally control the response in order to balance the risk of cancer with the risk of false responses to signals of DNA damage. To answer this question, we modularly study how the topology of alternative *p*53 core regulation networks integrate DNA damage information and determine the range of dynamic responses to DNA damage. Using a novel algorithm to solve for the equilibrium points of the different network models, we explore the qualitatively different types of switching behavior that the various network configurations can admit. This provides an insight into how the different core regulation genes play a functional role in determining the *p*53 response to DNA damage, and further reveals a novel mechanism by which ARF could play the role of an apoptosis enhancer [25].

We validate our model for the *p*53 network in vertebrates by using it to explain many recent experimental results in a way that previous models are unable to do. Specifically, our model predicts that for a moderate amount of damage, the *p*53 level will pulse, with the number of pulses proportional to the amount of damage. When the damage becomes too high, the *p*53 level elevates monotonically to a high level and remains there. This matches experiments on *p*53 in human breast cancer cells [10, 26, 27], and also more recent experiments on human U2-OS cells [13]. An important prediction made by our model in this regard is that the monotonic switch of *p*53 to a high value in response to a large amount of DNA damage is a result of a property of the core regulation network model and not due to temporal effects, ensuring that the mechanism is robust. Our model also predicts that the upstream transduction kinases are responsible for activating *p*53 pulses when damage is present and that pulses in kinase levels are highly coupled with *p*53, matching recent experimental observations [10, 28].

This work also elucidates how the *p*53 responses exhibited by different network structures relate to the potential costs and fitness benefits of different organisms. We show that the range of behaviors permitted by the vertebrate network is crucial in minimizing the frequency of Type 1 errors while eliminating Type 2 errors in human cells. We also show that the complexity of the network found in vertebrates allows for other distinct strategies for tumor suppression, as has been observed in elephants [29]. The alternative network configurations in primordial organisms however, cannot express the range of behaviors that are observed in and permitted by vertebrate cells.

## Taxonomic representation of regulatory genes and network configurations

Among clinical cancers, mutations in *p*53 are among the most commonly detected [4], and *p*53 is widely recognized as playing a key role in tumor suppression in multicellular organisms [7, 23]. The *p*53 family of genes is at least one billion years old [5], appearing in organisms as diverse as protists, cnidarians, and mammals. The tumor supression role of the *p*53 ancestral gene appears to have been preserved over a large time span; its function in cnidarians is to protect the germline gametes from DNA damage, and this function persists in arthropods, annelids, molluscs, and vertebrates [5].

In humans, *p*53 is part of a sophisticated network of hundreds of genes [20] that cooperate to mediate cell fate decisions such as the initiation of cell-cycle arrest, DNA repair, senescence or apoptosis, as soon as DNA damage is detected [21]. This network includes a core regulation network [23] consisting of the proteins *MDM*2 [30–32], *PTEN*, [33, 34], and *ARF* [35–37] among a host of other upstream, down-stream and intermediate species involved in sensing, transduction and regulation [21–23].

Recent studies have uncovered multiple aspects of how these core regulation proteins interact. It is well known that *p*53 transcriptionally activates the *MDM2* gene. *MDM2* in turn antagonizes *p*53 forming a negative feedback loop around *p*53 [30, 31, 38]. *p*53 also activates the *PTEN* gene, and *PTEN* proceeds to down-regulate *MDM*2 through a series of interactions [34, 39, 40], which forms a positive feedback loop around *p*53. *ARF* is known to cause the translocation and eventual degradation of *MDM2* [35, 36, 41], and it has been recently discovered that *ARF* can also mediate the degradation of *MDM2* [37], leading to a positive feedback loop around *MDM*2. The network configuration of these four genes will henceforth be referred to as the *full network* 𝒩_*υ*_, and is illustrated in Fig. 1. Much of our understanding of the core regulation network interaction and dynamics in response to DNA damage comes from studies of human and mouse cell lines [42], and this network is most likely common to at least mammals. As we will see in this paper, this network configuration is crucial in explaining experimental observations about *p*53 behavior in human cells in response to DNA damage.

**Fig. 1.**
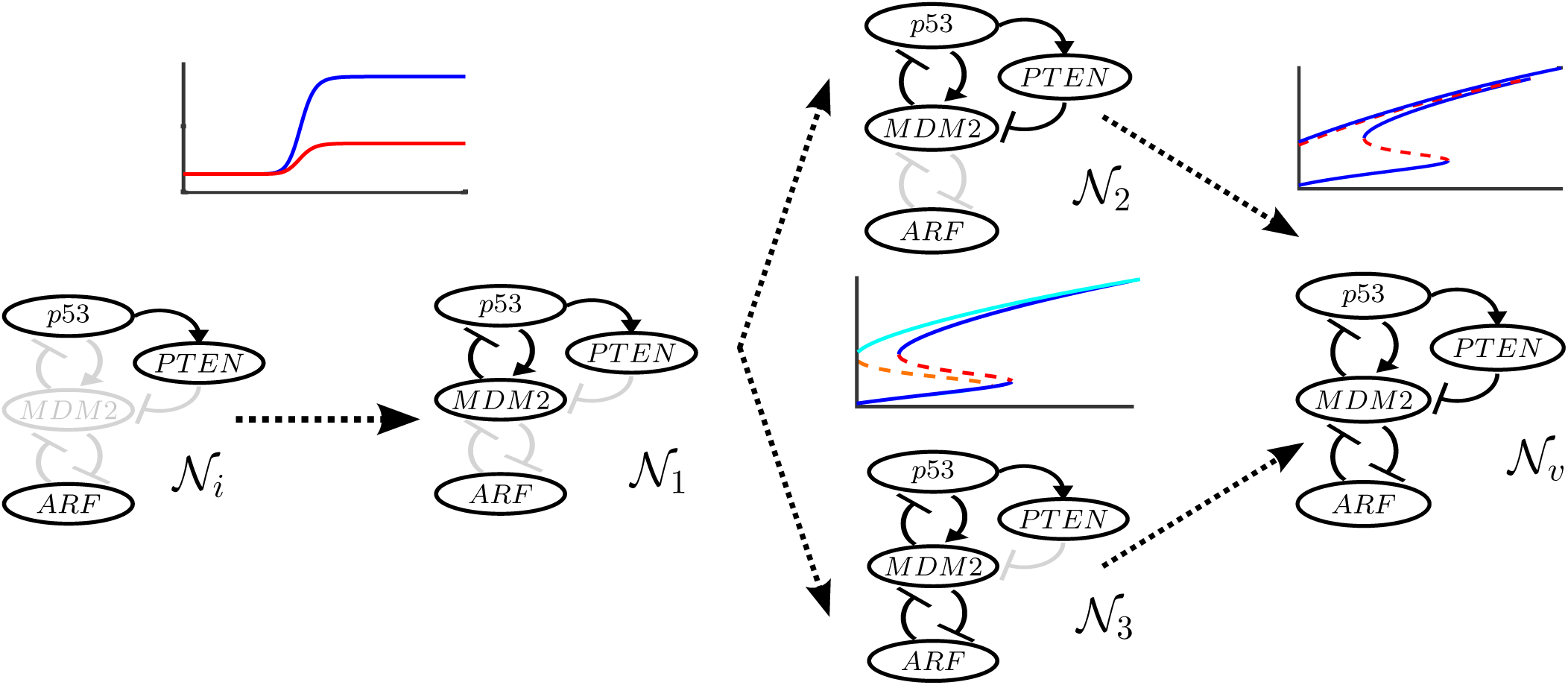
Evolutionary pathway to complexity from putative primordial organisms to modern-day vertebrates.

While homologs of *p*53, *PTEN* and *ARF* are present in a large taxonomic group of animals, including non-vertebrates, less evidence exists for the functional role of *MDM*2 outside of vertebrates. Recent work suggests that *MDM*2 and its paralog *MDM*4 arose from a gene duplication event and became the primary negative regulators of the *p*53 family of genes [43]. While invertebrate lineages contain a *MDM*2 homolog, it bears little functional resemblance to the vertebrate *MDM*2 and there is no direct evidence that it plays a role in regulating *p*53 dynamics. Examining the Network 𝒩_υ_ in Fig. 1 reveals that the absence of this *MDM*2 would prevent *PTEN* and *ARF* from affecting the dynamics of *p*53, because both *PTEN* and *ARF* only regulate *p*53 through feedback connections involving *MDM*2. This means that in the absence of *MDM*2, the response of *p*53 to DNA damage would only depend on the direct regulation of *p*53 by upstream inputs. This suggests that the dynamics of the network may depend critically on the feedback loops that *MDM*2 mediates and motivated us to explore the possible intermediate network configurations that might have emerged in the evolution from the putative primordial gene network to that observed in modern vertebrates.

### Evolutionary history of the p53 network

Based on the tax-onomic representation of *p*53 and *MDM*2, we hypothesize that the *p*53 function as a tumor suppressor evolved prior to the feedback loops mediated by *MDM*2. This hypothesized ancestral network is labeled 𝒩_*i*_ and is shown in Fig. 1. Following from the work of Momand et al [43], transitions to the network observed in modern mammals must have proceeded through the intermediate networks shown in Fig. 1. Two possible evolutionary paths are possible from the ancestral network 𝒩_*i*_ to the mammalian network 𝒩_υ_. In both evolutionary paths, the *p*53–*MDM*2 negative feedback interaction evolves first since this interaction is central to the regulation of *p*53 by either *PTEN* or *ARF*, leading to the network 𝒩_1_ shown in Fig. 1. In the first path, the interactions through which *p*53 inhibits *MDM*2 through *PTEN* evolves first (leading to the network 𝒩_2_ as seen in Fig. 1), followed by the mutually inhibitive feedback loop between *MDM*2 and *ARF*, leading to the network 𝒩_υ_. In the other path, the mutually inhibitive feedback loop between *MDM*2 and *ARF* evolves first, leading to the network 𝒩_3_. This is followed by the interactions through which *p*53 inhibits *MDM*2 through *PTEN*, leading again to the network 𝒩_υ_.

Our primary emphasis is on providing insight into the behaviors permitted by these different network configurations from a dynamical systems standpoint. As such, we do not make claims that the paths to complexity studied in this paper are the only possible routes that evolution might have evolved. Rather, we focus on exploring the different qualitative behaviors permitted by the intermediate network configurations and distinguish these from those observed in the mammalian network.

## Modular models of the *p*53 pathways

The diagrams shown in Fig. 1 are conceptual in that, while each block is labeled with the name of a single protein, it typically represents several interacting chemical species. For example, the block labeled *MDM*2 includes the dynamics of the *MDM*2 protein in its active an inactive forms, as well as its corresponding mRNA. We should thus think of each block as a “module” rather than an individual protein.

To predict how the network behavior changes as modules are added, the different modules in Fig. 1 must be associated with precise mathematical models, so as to exhibit two key properties: *dynamic modularity* and *parametric modularity.* Dynamic modularity ensures that the properties of each module do not change upon interconnection with other modules, and is therefore an essential property to ensure that the independent analyses of different modules can be combined to make precise statements about the behavior of the entire network, and specifically, how the behavior changes as modules are removed or inserted into the network. This can enable the analyses of large networks and also the design of novel networks [44–46]. Parametric modularity, which has not been explicitly mentioned as often in the literature, implies that the network parameters that are associated with a given module do not appear in any other module. This property ensures that each module can be analyzed independently to understand how various parameters or inputs to that module affect the outputs from the module.

Fig. 2 shows a mathematically precise “block-diagram” representation of the full network configuration from Fig. 1. Each module, labeled from 𝒨_1_ to 𝒨_4_, is characterized by a system of ordinary differential equations (ODEs) depicted in Fig. S1 that describe the behavior of the species in that module. The solid arrows between the modules represent the signals that flow between them. For example, the arrow originating at 𝒨_2_ and ending at 𝒨_1_ represents the mechanism by which active *MDM*2 represses *p*53. This signal is said to be an “output” of 𝒨_2_ and an “input” to 𝒨_1_. A more comprehensive block diagram representation of this network and the corresponding block diagram representations of the other network configurations are shown in Figs. S2 and S3 of the Supporting Information, respectively.

**Fig. 2.**
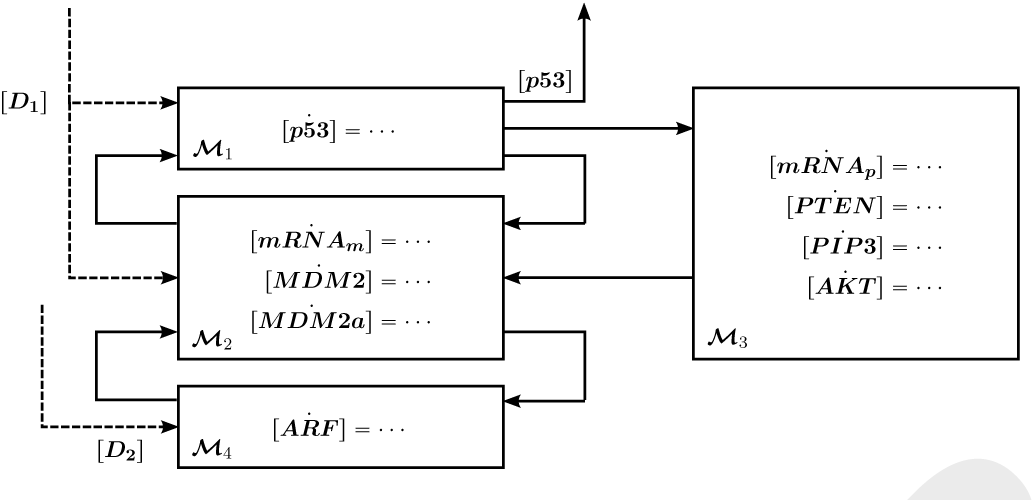
Modular block-diagram representations of full network 𝒩_υ_. The specific ODEs describing each module are shown in Fig. S1.

The species associated with each module are as follows. Module 𝒨_1_ is associated only with the dynamics of the *p*53 protein. Module 𝒨_2_ consists of the dynamics of *MDM*2 in both its active (*MDM2a*) and inactive (*MDM*2) forms, along with its mRNA (*mRNA_m_*). Module 𝒨_3_ consists of the dynamics of *PTEN* and its mRNA *(mRNA_p_),* along with the intermediate species *PIP*2 and its phosphorylated form *PIP*3, and *AKT* and its active phosphorylated form *AKTa.* Finally, Module 𝒨_4_ contains the dynamics of the *ARF* protein.

The inputs to the full network are *D*_1_, which represents kinases that activate *p*53 and cause the degradation of *MDM*_2_ in response to DNA damage [40], and *D*_2_, which represents transducers like *PARP*1 and *E2F*1 which activate *ARF* in response to DNA damage [25, 47]. A more detailed description of the model and its parameters is provided in the Supporting Information.

## The role of each *p*53 network configuration in determining switching behavior

In this section, we discuss the different dynamic behaviors that are permitted by each *p*53 network configuration we identified on the pathways to complexity. A specific type of behavior that we study is the ability of the *p*53 concentration to behave like a “switch” in response to DNA damage.

### Multistability and biological switches

Biological networks commonly use switch like behavior to turn a continuous input signal (usually an extrinsic stimuli) into an “all-or-nothing” output signal response (usually species or compound concentrations). A common mechanism employed by these networks to bring about this behavior is *bistability.* A bistable network has two sets of stable steady states, in which the state of the network transitions from a neighborhood of one set of steady states (which we call *A*) to the other (which we call *B*) when an input signal grows above a certain threshold (which we call *T_1_)*. A unique property of these networks is that once the transition occurs, even if the input later falls below *T*_1_, the state may still remain in a neighborhood of *B*. Reversion to a neighborhood of *A* is generally observed only when the input signal falls below another threshold *T*_0_, where *T*_0_ < *T*_1_. This property is known as hysteresis, and refers to the ability of a bistable system to “remember” that the input stimulus was above *T*_1_ long after that stimulus is removed, until it falls below *T*_0_ [48]. The lac operon network in E. Coli, the Weel Cdc2 network, the *NF_kβ_* response in an apoptosis network and the cell cycle oscillator in *Xenopus laevis* are among the many naturally occurring biological networks whose responses have been modeled as bistable switches [48–50]. In some cases, the lower threshold *T*_0_ does not exist, which makes the transition from *A* to *B irreversible*. It is also plausible to have more than two sets of stable steady-states, which is known as *multi-stability*.

*Monostable networks* on the other hand have states that slide along a continuum of steady-states [49] in response to input signaling. Such networks can only exhibit switch-like behavior if the system is *ultrasenstive;* that is, if the inputoutput relationship of the network is described by a sharp sigmoidal curve [48]. When the input signal crosses the exponential phase of the sigmoid, the output response of the network transitions from a low to a high state. In fact, the dynamics of transcription factor target genes as a function of transcription factor concentration is typically modeled as a sigmoidal “Hill function”, which can be viewed as an ultrasensitive switch. However, such switches are *memoryless:* once the input signal is removed or is reduced, the system immediately returns to its original state [48].

The different *p*53 network configurations can display various switching mechanisms with respect to the two damage sensing inputs [*D*_1_] and [*D*_2_]. Our modular study revealed that as [*D*_1_] is varied, the network configurations 𝒩_*i*_ and 𝒩_1_ operate as monostable switches, in the sense that when an increase in [*D*_1_] makes the *p*53 level rise, a subsequent decrease in [*D*_1_] to its original value causes the concentration of *p*53 to revert to its initial level. The steady-state concentration of *p*53 as a function of [*D*_1_] is shown in Fig. 3. The main difference between the two networks 𝒩_*i*_ and 𝒩_1_ is that negative feedback of *p*53 with *MDM*2 typically makes the *p*53 steady-state in 𝒩_1_ to be lower than that of 𝒩_*i*_.

**Fig. 3.**
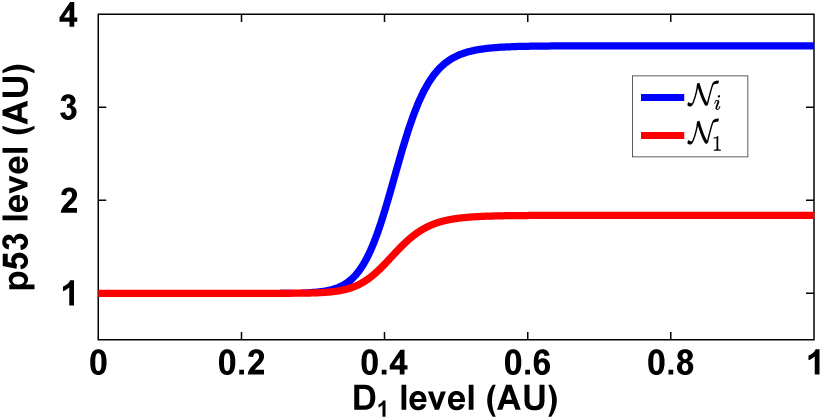
Steady-state concentration of *p*53 as a function of [*D*_1_] shows a continuum of steady-states in both 𝒩_*i*_ and 𝒩_1_. Due to the negative feedback on *p*53, 𝒩_1_ admits lower *p*53 steady-state values than 𝒩_*i*_. The *p*53 levels are normalized by their value when [*D*1] = 0.

### Networks with a positive feedback loop admit switching with “memory”, allowing for sustained high levels of *p*53 expression once damage crosses a threshold

With model parameters similar to those from [40], the network 𝒩_2_ operates as a bistable switch with respect to the input [*D*_1_]. From this study, we can plot the *bifurcation diagram* of 𝒩_3_ with respect to the parameter [*D*_1_] in Fig. 4(a). The solid blue line signifies a continuum of stable steady-state concentrations of *p*53 as a function of [*D*_1_], while the red dashed lines represent unstable equilibrium points. It can be seen that when [*D*_1_] is between the two thresholds *T*_0_ and *T*_1_, there are two stable steady-states the *p*53 level can converge to. The bifurcation diagram and simulations are normalized by the *p*53 level at [*D*_1_] = *T*_0_ for clarity.

**Fig. 4.**
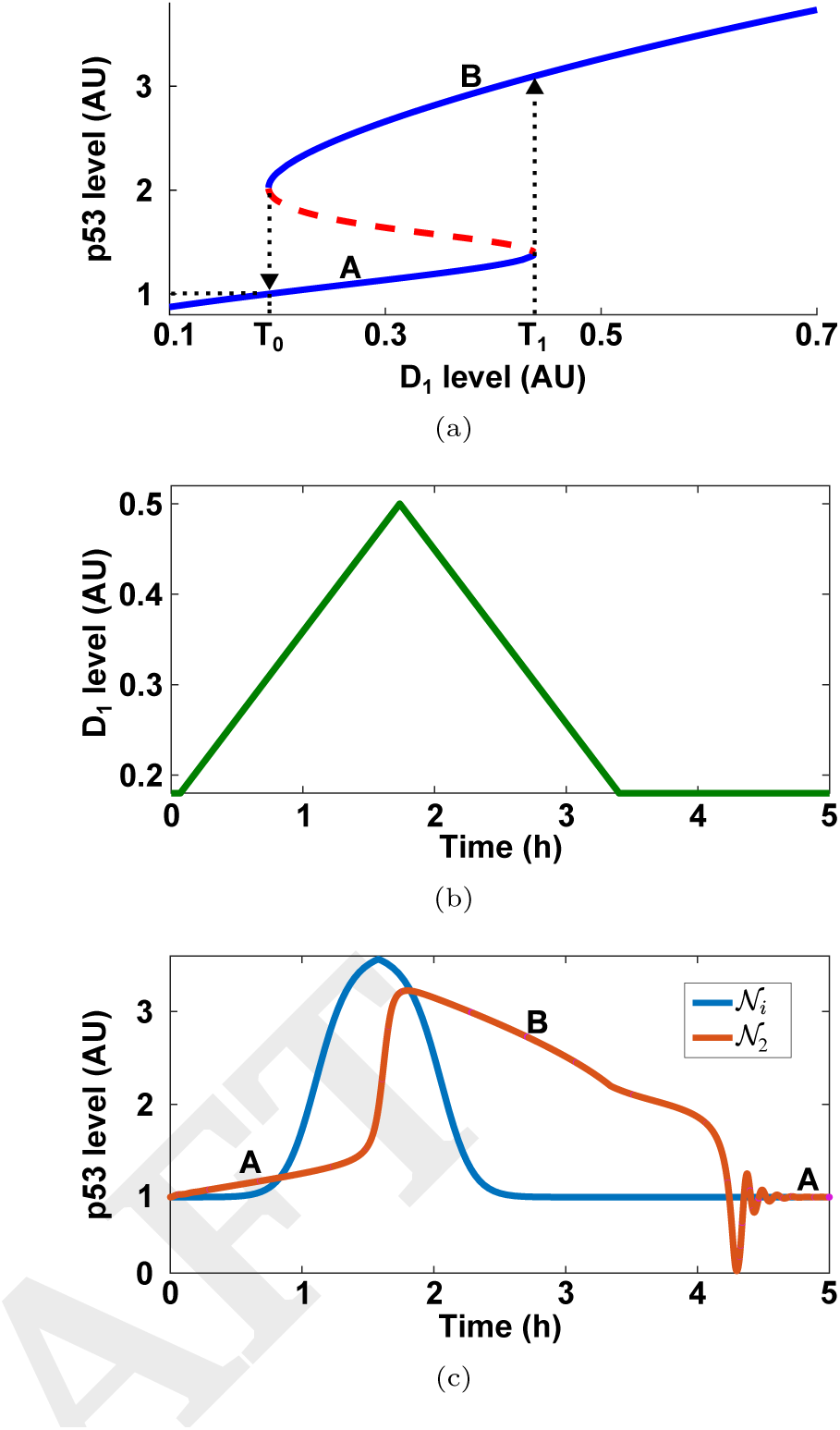
(a) Bifurcation diagram of 𝒩_2_, illustrating the reversible bistable switch. Assuming the *p*53 level starts at a point in “A”, increasing [*D*_1_] beyond *T*_1_ results in the *p*53 level making a switch to a point in “B”. For the *p*53 level to revert to a point in “A”, the [*D*_1_] needs to fall below *T*_0_. (b) The level of *D*_1_ in our simulation. (c) The corresponding *p*53 response contrasting the behaviors of the monostable network 𝒩*i* to the bistable network 𝒩_2_ as [*D*_1_] is varied. While the bistable switch has memory, the monostable switch does not.

To illustrate the behavior of the bistable network, we run a simulation as shown in the red curve of Fig. 4(c). We observe the manifestation of the “memory” property of bistable networks, where after rising to a point in set of equilibrium points labeled as “B”, the *p*53 level returns to a point in the set “A” after [*D*_1_] crosses the threshold *T*_0_ from above. Fig. 4(c) further contrasts this to the behavior of the monostable networks 𝒩_*i*_ (blue curve), where the switch has no memory.

A change in the parameters corresponding to the rate of initiation of *p*53 and ubiquitination of *MDM*2 by *D*_1_ causes the threshold *T*_0_ from Fig. 4(a) to become less than 0, as shown in Fig. 5(a). In this case, the network exhibits an irreversible switch from the set of points in Region “A” to those in Region “B”. This is illustrated in Figs. 5(b)–(c). The bifurcation diagram and simulations are normalized by the *p*53 level at [*D*_1_] = 0.2 for clarity.

**Fig. 5.**
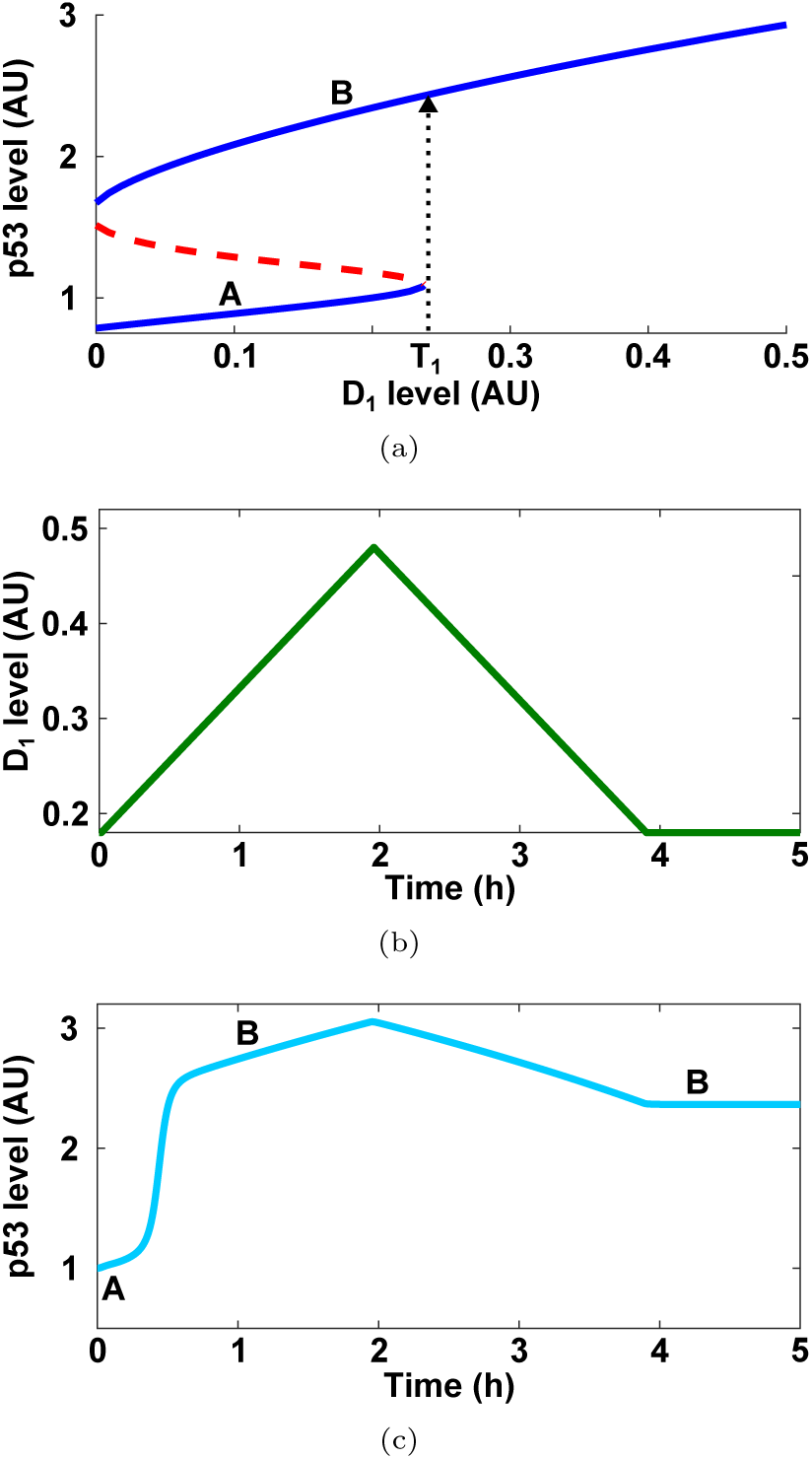
(a) Bifurcation diagram of 𝒩_2_, illustrating the irreversible bistable switch. (b) The level of *D*_1_ in our simulation.(c) The corresponding response of *p*53 demonstrating the irreversible switch.

The network 𝒩_3_ can also behave as a bistable switch with respect to [*D*_1_] provided that [*D*_2_] also increases along with [*D*_1_]. This is a reasonable assumption, since both [*D*_1_] and [*D*_2_] will increase with DNA damage. Qualitatively, this network admits the same behavior as 𝒩_2_. The bifurcation diagram for this network is provided in Fig. S4 of the Supporting Information.

Unlike in the monostable networks, bistability allows for the *p*53 levels to remain high for a sustained period of time once the damage sensor level has crossed threshold *T*_1_, even if the damage sensor level subsequently falls below this level. This can induce *p*53 to up-regulate down-stream pathways that bring about cell-cycle arrest and DNA repair, as well as those that bring about apoptosis or senescence. Since apoptotic pathways are typically slower than repair pathways [51], it is more likely that apoptosis will be the cell fate either if the damage sensor level remains high for sufficiently long, or if the network operates as an irreversible bistable switch, since a sustained high levels of *p*53 expression is known to bring about apoptosis [12, 16, 18].

### The full network 𝒩_υ_ can admit both a reversible and an irreversible switch, allowing for distinct *p*53 responses to different levels of DNA damage

The addition of the *ARF* module 𝒨_4_ to 𝒩_2_, or the *PTEN* module 𝒨_3_ to 𝒩_3_, can bring about *tri-stability* in the *p*53 levels with respect to [*D*_1_]. That is, there are three sets of stable steady-states between which the *p*53 level can toggle, which we label “A”, “B” and “C”.

The bifurcation diagram illustrating the steady-states of *p*53 in this network 𝒩_υ_ is shown in Fig. 6(a). From this diagram, we observe that the original switching behavior which was a property of the bistable networks is retained; when [*D*_1_] crosses *T*_1_ the *p*53 level switches to a high value, and this switch is “turned-off” only when [*D*_1_] falls below *T*_0_. In addition to this behavior, the tri-stable network admits a third threshold *T*_2_, that can be seen in Fig. 6(a). Once [*D*_1_] crosses *T*_2_, *p*53 transitions *irreversibly* to a high value; no matter how [*D*_1_] changes following this transition, *p*53 will not return to its nominal level. This behavior is illustrated in Figs. 6(b)–(c). The bifurcation diagram and simulations are normalized by the *p*53 level at [*D*_1_] = *T*_0_ for clarity.

**Fig. 6.**
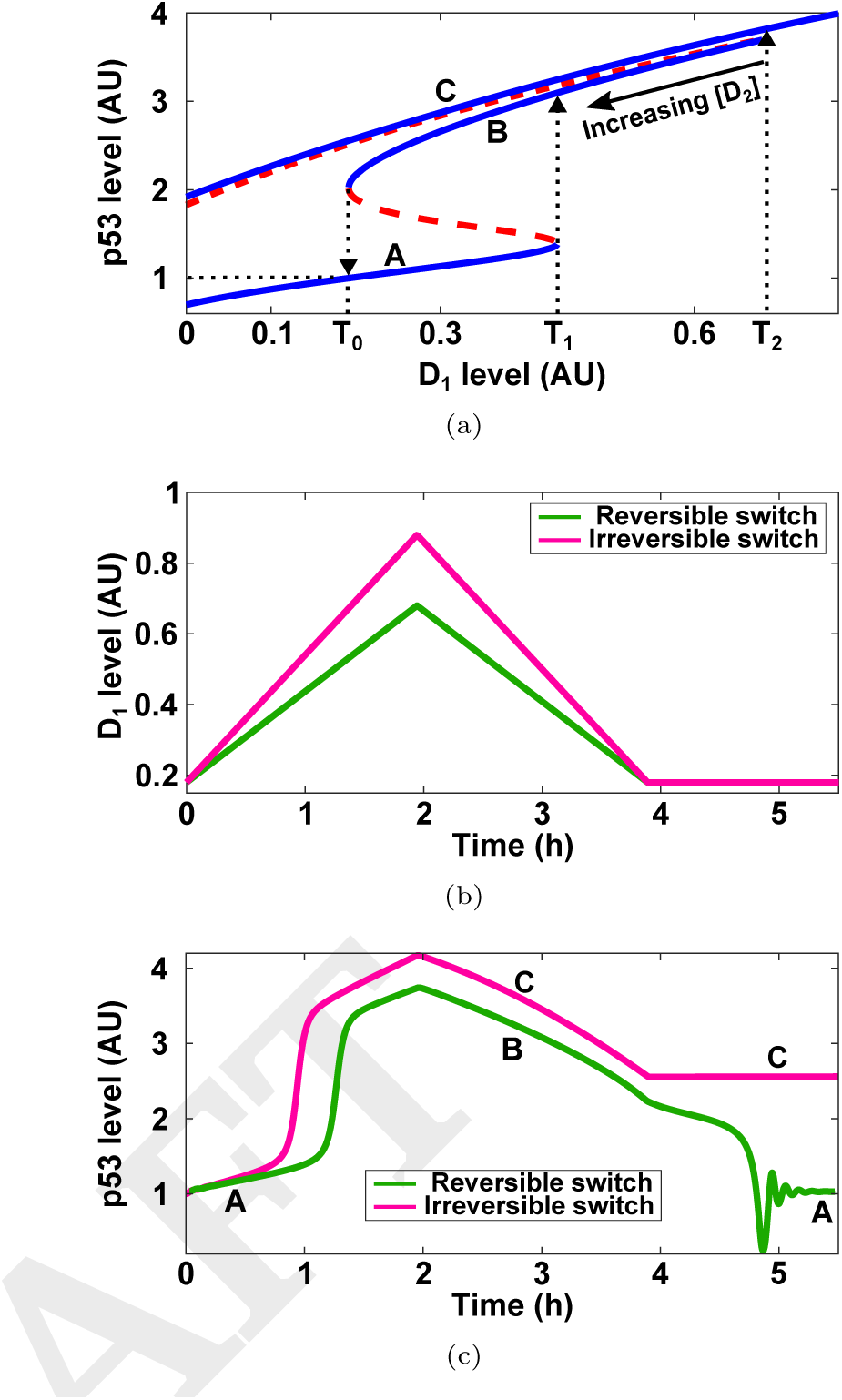
(a) Bifurcation diagram of the network 𝒩_υ_, illustrating the tristable switch. The switching behavior of the bistable network is retained. However, there is an additional threshold *T*_2_ which when [*D*_1_] crosses, results in an irreversible switch of the *p*53 level to the set of points “C”. (b) Simulated *D*_1_ levels. [*D*_2_] is kept fixed. (c) The resulting *p*53 behavior shows that the tristable network can behave both as a reversible and an irreversible switch for different values of maximal [*D*_1_]. The green line represents the reversible switch, while the pink line represents the irreversible switch.

While the bistable networks can operate as either reversible or irreversible switches, the full network is able to simultaneously admit both switching behaviors. From the perspective of our network, the level of damage-sensor [*D*_1_] crossing *T*_1_ is likely associated with a scenario that enough damage has been triggered to initiate a *p*53 repair response, but not to bring about a terminal cell fate. On the other hand, [*D*_1_] crossing *T*_2_ likely means that too much damage has been sustained, causing an irreversible switch in the *p*53 level to a high value and eventually bringing about either senescence or apoptosis.

It turns out that the level of *D*_2_ also plays an important role in shaping the *p*53 response. We recall that *D*_2_ represents transducers like *PARP*1 and *E2F*1 which activate *ARF* in response to DNA damage. As [*D*_2_] increases, the rate of pro-duction of *ARF* is increases. This in turn causes the middle branch of stable equilibrium points (labeled ‘B’) to “shrink”, as illustrated in Fig. 6(a). A lower threshold *T*_2_ would mean that the cell has a higher chance of going to senescence or apoptosis, since the irreversible switch in *p*53 levels is triggered for lower levels of [*D*_1_]. This behavior illustrates how *ARF* activation can trigger apoptosis, as has been observed experimentally [25]. The dynamical behaviors permitted by the different networks are summarized in Fig. 1, which also shows the possible evolutionary paths from putative primordial organisms to extant vertebrates.

## Simple models of DNA damage transduction and repair explain experimentally observed results

We now show how our tri-stable model for the *p*53 core regulation network in vertebrates 𝒩_υ_, coupled with simple models of DNA damage transduction and repair, explains many experimentally observed results. We model the activity of the transduction kinases such as *ATM* and *CHK1* using a single variable *D*_1_, similar to what was done in [10]. We model the dynamics of *D*_1_ as

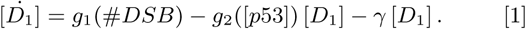

In this equation, the variable *#DSB* represents the number of DNA double-strand breaks (DSBs), which activate *D_1_* through the function *g_1_(#DSB*). *D*_1_ then goes on to activate *p*53 and cause the accelerated degradation of *MDM2* [32, 40], per the model in Fig. 2. The feedback term *g*_2_([*p*53]) represents negative feedback between *p*53 and the transduction kinases [10, 28], and the rate constant *γ* represents the *p*53-independent degradation of the kinases. We assume that the dynamics of Eq. (1) is slow compared to the dynamics of the *p*53 core regulation network, which is a key element in explaining the experimentally observed behavior.

We model the dynamics of *#DSB* as a simple deterministic death process,

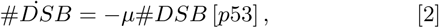

where *μ#DSB* [*p*53] is the rate of repair, that is known to be regulated by *p*53 [52–54]. Since damage is induced in the cell on the order of minutes [55], while the response can last for many hours or even days [26, 27], the number of DSBs induced by damaging agents is introduced as the initial condition of Eq. (2).

*D*_2_ represents the activity of proteins such as *PARP*1 [47] and *E2F*1 [25], which are known to play a role in activating *ARF* in response to DNA damage. We model [*D*_2_] as a simple monotonically increasing function of the number of DNA DSBs

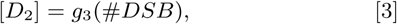

with the rate of production of *ARF* governed by [*D*_2_]. The specific parameters and functions from Eqs. (1)–(3) are given in the Supplemental Information.

We couple these models with the full network 𝒩_υ_ and report on the resulting simulations. In Fig. 7(a), we show that the *p*53 levels remain low for a damage resulting in 5 DNA DSBs, which is a low level of DNA damage. In this case, the DNA damage is repaired before a *p*53 pulse is initiated. For damage resulting in 10 DSBs, a single *p*53 pulse is observed as shown in Fig. 7(b). This matches experimental results from [26], which show that O.3G*_γ_* of damage is sufficient to initiate a pulse, where 1G*_γ_* of damage results in between 30-35 DSBs [27, 56, 57]. In Fig. 7(c), our simulation shows that DNA damage that brings about 150 DSBs causes the initiation of multiple *p*53 pulses for over 60 hours until the damage is repaired, matching experimental results that suggest that the number of pulses increases as the number of DSBs increases [4, 26, 27], and that these pulses could last for days after damage is induced [27]. All pulses in our simulation have a period of between 4-7 hours, matching these experimental results in human breast cancer cells.

**Fig. 7.**
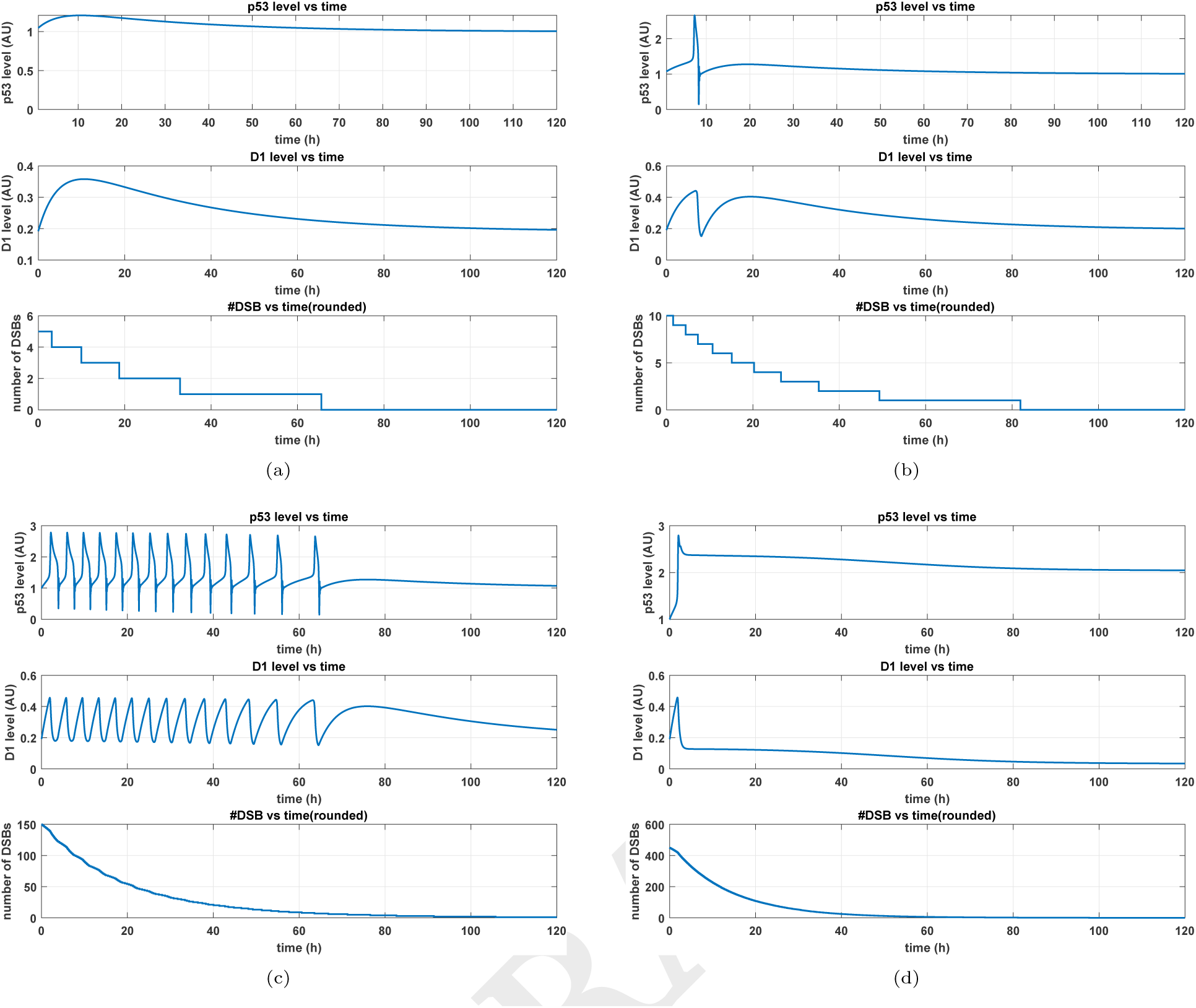
*p*53 and *D*_1_ responses, along with DSB repair process for (a) Initial #*DSB* = 5 in 𝒩_υ_ (b) Initial *#DSB* = 10 in 𝒩_υ_ (c) Initial *#DSB* = 150 in 𝒩_υ_ (d) Initial *##DSB*## = 450 in 𝒩_υ_

Early modeling work suggested that these pulses were primarily a result of the negative feedback interaction between *p*53 and *MDM*2, along with some positive feedback or delays [55, 58–61]. However, more recent experimental results, in which pulses in *p*53 are observed to be highly coupled with pulses in the upstream transduction kinases, suggest that *p*53 pulses are brought about by the interaction of *p*53 with these kinases [10, 28]. Another important observation from [10] is that transient activation of the kinases results in a pulse of *p*53, suggesting that there is an excitable mechanism controlling *p*53 pulses [10, 28]. It was noted that while a single negative feedback loop between *p*53 and these kinases (with delays) would be sufficient to generate sustained oscillations, other *p*53 interactions would be necessary to explain the observed excitability.

In our model, the interaction between *D*_1_ and the *p*53 network 𝒩_υ_ can be described as an excitable mechanism known as a relaxation oscillator [62]. The slow negative feedback between *D*_1_ and *p*53 described in Eq. (1) operates over the faster dynamics of the *p*53 core regulation network. The pulses in *p*53 are observed because of the resulting cyclic transfers between the *p*53 equilibrium points in Regions ‘*A*’ and ‘*B*’ [Fig. 6(a)]. The pulses in *D*_1_ levels are also seen to be highly coupled with the pulses in *p*53 levels. In this way, our simulation results strongly matches both experimental observations, and hypotheses about the underlying mechanism that brings about the observed behavior without recourse to additional mechanisms or interactions.

In Fig. 7(d), our simulation shows the *p*53 levels monotonically elevating to a high level for a large number of DNA DSBs, and remain high without pulsing. Even with the progression of the DNA repair process, the *p*53 levels remain high, due to the irreversible switch permitted by the tristable model. This behavior also matches recent experimental observations in human U2-OS cells, in which the *p*53 pulsing is observed for moderate levels of DNA damage, but very high levels of DNA damage trigger a strong monotonic elevation of the *p*53 level that does not return to a low level after rising [13].

It is worth noting that the experimental results are from different cells. Our goal is to show that the experimentally observed behavior is possible, and not to identify specific parameter values for the different cell types.

## Discussion

Some of the earlier *p*53 modeling work was carried out under the assumption that the transduction kinase levels were proportional to the amount of damage, and that pulses were primarily observed due to the negative feedback between *p*53 and *MDM*2 [58, 59]. Other modeling work, while considering the positive feedback loop, assumed that the upstream kinase concentrations were proportional to damage and caused autonomous pulsing in *p*53 when the kinase levels were sufficiently high [55, 60, 61, 63]. In contrast, our model takes into account recent experimental results which suggest that upstream transduction kinases are responsible for initiating *p*53 pulses, and that the pulses of these kinases are highly coupled with pulses of *p*53 [10, 28]. Moreover, these previous models primarily explain plausible mechanisms for the observation of oscillations in *p*53, and do not postulate the irreversible switch in the concentration of *p*53.

There are other models of the *p*53 core regulation network in humans that display both the pulsing and sustained high levels of *p*53 [40, 63, 64]. However, our model matches recent experimental observations that demonstrate pulsing in *p*53 for a moderate amount of damage, and a strong monotonic elevation in *p*53 for high levels of damage [13], contrasting to these previous models in which *p*53 pulses before switching to a high value [40, 63, 64].

To our knowledge, our model of *p*53 core regulation in vertebrates is the first where the irreversible switch of *p*53 to a high level is an inherent property of the tristable network structure, as opposed to a consequence of temporal behavior. It is also worth noting that all the observed behavior can be explained precisely using the dynamical properties of the network ODE model, and do not necessitate the artificial introduction of external signals, switches or clamps.

A novel behavior that our model predicts is the role played by *ARF* in bringing about an irreversible cell fate. It has long been postulated that *ARF* enhances apoptosis by sequestering away *MDM*2, thereby causing an increase in *p*53 levels. Although models to capture this behavior have been proposed [35, 65], the details of the underlying dynamics are still largely unknown [66]. A recent study has further pointed to a novel mechanism through which *MDM*2 can inhibit *ARF* [37]. In our model, this mutually inhibitory feedback between active *MDM*2 and *ARF* is crucial in bringing about tristability. Our model predicts that an increased rate of production of *ARF* is an important factor in bringing about the irreversible switch of *p*53 to a high level. In this way, our model proposes a novel and mathematically sound reasoning for how *ARF* might behave as an apoptosis enhancer.

From an evolutionary standpoint, we can understand the selection pressures on the tumor suppressor network as balancing a trade-off between minimizing the number of Type 1 errors, while eradicating Type 2 errors. Type 1 errors occur when the statistical null hypothesis that DNA can be repaired to its original state is true, but the cells are killed anyway, while Type 2 errors occur when the null hypothesis is false, but the cells are not killed. While Type 1 errors could have metabolic costs in that too many cells are killed, Type 2 errors are potentially fatal.

From this perspective, the functional role of the *p*53 core regulation network in humans is clear. For low amounts of damage, the *p*53 level remains low. This is associated with the fact that the damage is deemed too low for a tumor to develop, and hence a significant *p*53 response is not initiated.For moderate amounts of damage, *p*53 exhibits pulsatile behavior, with the number of pulses proportional to the amount of damage. At this point, the damage triggers the repair mechanisms by inducing pulses in *p*53 until all the damage has been repaired. This is consistent with experimental results which show that majority of cells that exhibit pulsed *p*53 are able to grow and divide after recovery from DNA damage [12]. For a sufficiently large amount of damage, the *p*53 level switches monotonically to a high level and remains high. In cells that exhibit sustained high levels of *p*53 signaling, most cells go into senescence or apoptosis [12, 13, 18]. This switch to fixed high *p*53 levels removes the possibility that a reduction in the damage signal could allow unrepaired cells to proliferate. In this way, the cell is able to minimize the number of Type 1 errors by repairing DNA damage for a range of moderate levels of damage, but also has a clear threshold to bring about an irreversible switch to apoptosis when the amount of damage sustained is too high.

If every cell had an equal probability of developing cancer per unit time across all organisms, then large long-lived
mammals would have a very high risk of developing cancer as compared to smaller short-lived organisms. However, there is a lack of correlation between the body size of an organism and its cancer risk [67], and this is known as “Peto’s Paradox”. There are many hypotheses to resolve Peto’s Paradox, as outlined in [67]. Two of these hypotheses, having a redundancy of tumor suppressor genes and a more sensitive apoptotic process, were found to be true in a recent study on elephants [29]. It was found that elephant cells have 20 copies of the *TP*53 gene as compared to the single copy observed in human cells. Elephant cells were also found to carry out *p*53-dependent apoptosis at a much higher rate than human cells for the same amount of DNA damage [29].

To test the effect of increasing the number of copies of the *TP*53 gene, we took the full vertebrate network 𝒩_υ_ described above and increased the basal and kinase-dependent rates of production of *p*53, while keeping all other parameters constant. We found that this would cause the level of threshold *T*_2_, shown in Fig. 6(a), to decrease. This change results in *p*53 making an irreversible switch to a high level and hence bring about apoptosis for a smaller number of DSBs, which would explain the experimental results observed in elephants [29]. A key point to note is that the increased apototic rates result from the temporal dynamics of the network, rather than from a simple increase in the basal concentration of *p*53. Our model also predicts that increasing the *ARF* sensitivity to DNA damage would also have a similar effect, and it would be interesting to study this gene in other large long-lived mammals.

The networks 𝒩_2_ and 𝒩_3_ were found to admit both reversible and irreversible bistable switching behavior. It is worth noting that these networks are not capable of exhibiting tristability regardless of the values of the model parameters.

For the reversible switch shown in Fig. 4(a), the *p*53 dynamics of these networks would be qualitatively similar to that of 𝒩_υ_ in the presence of low and moderate amounts of DNA damage when coupled with the transducer and repair dynamics we introduced. When the DNA damage is very high, the *p*53 level monotonically elevates to a high level. However, this switch is reversible if the DNA damage is repaired before the cell goes into senescence or apoptosis. This is illustrated for 𝒩_2_ in Fig. S5 of the Supporting Information. However, since the cell would have experienced a large amount of damage, the DNA repair may not have been successful or may have introduced genome destabilizing mutations. Hence, while the reversible bistable switches can minimize Type 1 errors by re-pairing moderate levels of damage, the elimination of Type 2 errors in the bistable networks requires the fast initiation of apoptotic pathways once the *p*53 level becomes high.

In the case of the irreversible switch shown in Fig. 5(a), *p*53 switches irreversibly to a high level once [*D*_1_] crosses threshold *T*_1_. However, in this case *p*53 is not able to admit pulsatile behavior, increasing the chance of the cell going to senescence or apoptosis. Hence, this policy would be very effective in eliminating Type 2 errors, but could also increase the frequency of Type 1 errors. In essence, the bistable networks 𝒩_2_ and 𝒩_3_ are either able to admit *p*53 pulses or an irreversible switch of *p*53 to a high level triggering apoptosis, but not both phases of behavior as the tristable network is able to. These networks would probably have been more likely to behave as irreversible switches, since the elimination of Type 2 errors is crucial in the cell’s ability to avoid tumors.

The network configurations 𝒩_*i*_ and 𝒩_1_ will only admit a single continuum of stable equilibrium points [68], and hence can only exhibit ultrasensitive switching behavior. In the absence of more complex mechanisms, the *p*53 levels are incapable of pulsing. Moreover, when the *p*53 level is high, it is more sensitive to changes in *D*1 levels than the bistable switch. In these organisms, it is therefore likely that the exponential phase of switching happens for reasonably low amounts of damage, and that the pro-apoptotic pathways were very sensitive to rising levels of *p*53, to ensure that Type 2 errors do not occur. This hypothesis is consistent with an earlier proposal on the operation of the *p*53 family of genes in early organisms [5].

In conclusion, our model shows that one adaptive value of the *p*53 core regulation network in vertebrates is its ability to balance the risk of cancer with the cost of too much cell death, and this behavior is clearly observed in experiments on human cells. The complexity of the *p*53 network in vertebrates also permits other organisms to operate at different points on this trade-off curve. On the other hand, inferred ancestral organisms with alternative *p*53 network configurations admit a narrower range of behaviors, which in turn limits the range of the possible points which they can lie on this trade-off curve.

## Materials and Methods

The computation of the bifurcation diagrams in Figs. 4(a), 5(a) and 6(a) requires the computation of equilibrium points for the system of ODEs in the corresponding networks. In practice, this reduces to the problem of computing the solutions to multivariate polynomial equations, which is known to be an NP-complete problem [69]. While simple equations can often be solved using commercially available numerical solvers, more complicated systems such as the ones that arise in our analysis, exhibit multiple equations with many possible solutions that are close to each other. In this case, solvers or numerical continuation software for the computation of bifurcation diagrams often miss solutions that might exist, which was the case when attempting to evaluate the equilibrium points of the tri-stable and bistable networks.

To overcome this problem, we developed an algorithm to solve for the equilibrium points of a network as a function of the inputs to the network. This algorithm was used to solve for the *p*53 equilibrium points as a function of the concentrations of the transducer inputs *D*_1_ and *D*_2_, to produce the plots in Figs. 4(a), 5(a) and 6(a).

A key aspect in the operation of the algorithm is the modular decomposition of the *p*53 network into the modules shown in Fig. 2. The procedure that the algorithm employs involves *gridding* the equilibrium values of the input and output signals for each module to form a table known as the *Steady-State Behavior* (SSB) of a module [70]. Typically, this table is obtained by gridding the equilibrium values of the outputs for a range of constant inputs, and this special case of the SSB has been termed the *Static Characteristic Function* (SCF) [71, 72] of the module. It is worth noting that these tables could be multi-valued, as there could be multiple output equilibria for a given input as is evidenced in the bistable and tri-stable networks.

Once the SSB of each module has been determined, the algorithm proceeds to systematically eliminate the *latent variables* in each of these tables, which are typically the input and output signals from each module that are interconnected to other modules in the network. This procedure is outlined in detail in the Supporting Information, and is crucial to obtaining and understanding the bifurcation behavior that each network configuration can admit.

## ACKNOWLEDGMENTS

This material is based upon work supported by the National Science Foundation under Grants No. ECCS-0835847 and EF-1137835. The authors would like to thank Bjørn Østman for his assistance with the evolutionary analysis.

## A Core regulation network models

In this section, we provide a detailed description of the model for *p*53 core regulation used in our work, along with the parameters used in simulations. This section describes the modules 𝒨_1_ to 𝒨_4_ that appear in Fig. 2 of the main text.

### A.1 Model description

The ordinary differential equations (ODEs) describing the dynamics of module 𝒨_1_ are depicted in Fig. 1(a). In this model, *p*53 is activated at some basal rate *p*_1_ and also in an input-dependent manner as a function of *u*_11_ = [*D*_1_], which represents kinases that activate *p*53 in response to DNA damage. *p*53 has a basal degradation rate constant of *d*_1_. The degradation of *p*53 is also mediated by the input *u*_12_, which typically describes the antagonism of *p*53 by active *MDM*2 [1, 2]. This results in the interconnection between Modules 𝒨_1_ and 𝒨_2_ given by

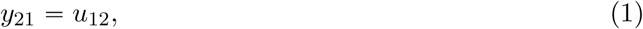

where *y*_21_ is an output from Module 𝒨_2_ given by the expression

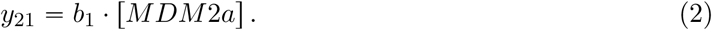

The ODEs describing the dynamics of module 𝒨_2_ are depicted in Fig. 1(b). In this model, *mRNA_m_* has a basal transcription rate *p*_2_ and a degradation rate constant *d*_2_. In addition, *mRNA_m_* transcription is activated by *p*53 through the input *u*_21_, which can be described through the interconnection

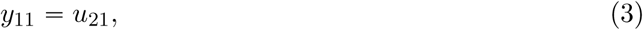

where *y*_11_ is an output from Module 𝒨_1_ given by the expression

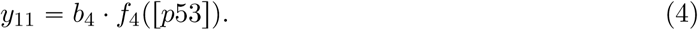

*MDM*2 is translated at a rate of *p*_3_ [*mRNA_m_*] and degraded with rate constant *d*_3_. The phosphorylation of *MDM*2 into its active form *MDM2a* is mediated by an input *u*_22_, which is typically mediated by *AKTa* that is associated with 𝒨_3_ [3, 4]. This results in the interconnection

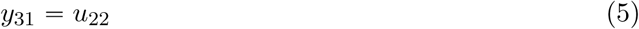

where *y*_31_ is the only output from Module 𝒨_3_ given by the expression

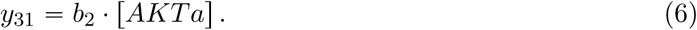

The rate of dephosphorylation back to *MDM*2 is given by *b*_3_ · *f*_3_{[MDM2a]). *MDM2a* has a basal degradation rate constant *p*_4_. The degradation of *MDM2a* is also mediated by two inputs *u*_23_ and *u*_24_. *u*_23_ = [*D*_1_] represents the concentration of upstream transduction kinases like *ATM*, which mediate the down-regulation of active *MDM*2 [5–7]. *u*_24_ represents the *ARF*-mediated translocation and eventual degradation of active *MDM*2 by ARF [8–1], which results in the interconnection

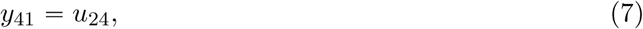

where *y*_41_ is the only output from Module 𝒨_4_ given by the expression

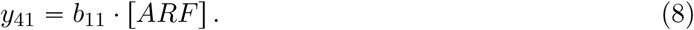

The ODEs describing the dynamics of the module 𝒨_3_ are depicted in Fig. 1(c). In this model, the mRNA of *PTEN*, represented by *mRNA_p_*, is transcribed at some basal rate *p*_5_, with a degradation rate constant *d*_4_. The transcription of *mRNA_p_* is also governed by the input *u*_31_, which describes the transcriptional activation of *PTEN* by *p*53 [4, 6, 11] resulting in the interconnection

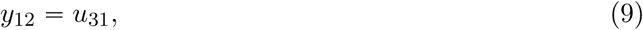

where *y*_12_ is an output from Module 𝒨_1_ given by the expression

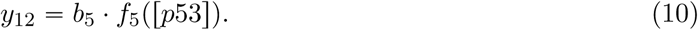

*mRNA*_*p*_ is translated to *PTEN* with rate constant *p*_6_, and degraded with rate constant *d*_5_. *PTEN* mediates the dephosphorylation of *PIP*3 into *PIP*2. The total concentration of *PIP2* and *PIP*3 is given by the constant *tot*_1_. The rate of phosphorylation of *PIP*2 is given by the term *b_6_f_6_{tot_1_ — [PIP*3]), while the rate of dephosphorylation of *PIP*3 is given by *b*_7_[*PTEN*]*f*_7_{[*PIP*3]). *PIP*3 then mediates the phosphorylation of *AKT* to *AKT a* (active *AKT*). The total concentration of *AKT* and *AKTa* is given by the constant *tot*_2_. The rate of phosphorylation of *AKT* is given by the term *b*_9_[*PIP*3] *f*_9_{*tot*_2_ — [*AKTa*]), while the rate of dephosphorylation of *AKTa* is given by bs *f*8,7{[*AKTa*]) [6, 12].

The ODEs describing the dynamics of module 𝒨_4_ are depicted in Fig. 1(d). In this model, *p*_7_ is the basal rate of production of *ARF*, with *u*_41_ = [*D*_2_] the extrinsic rate of production that represents the activation of *ARF* in response to DNA damage through transducers like *PARP*1 [13] and *E2F*1 [14]. *d*_6_ is the rate constant for the degradation of *ARF*. *ARF* is also degraded by an extrinsic process mediated by *u*_42_, which represents a novel recently-discovered mechanism through which *MDM*2 can mediate the degradation of *ARF* [15]. This results in the interconnection

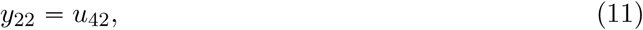

where *y*_22_ is an output of Module 𝒨_2_ given by the expression

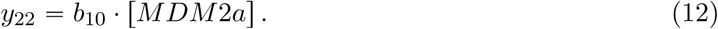

**Figure S1:**
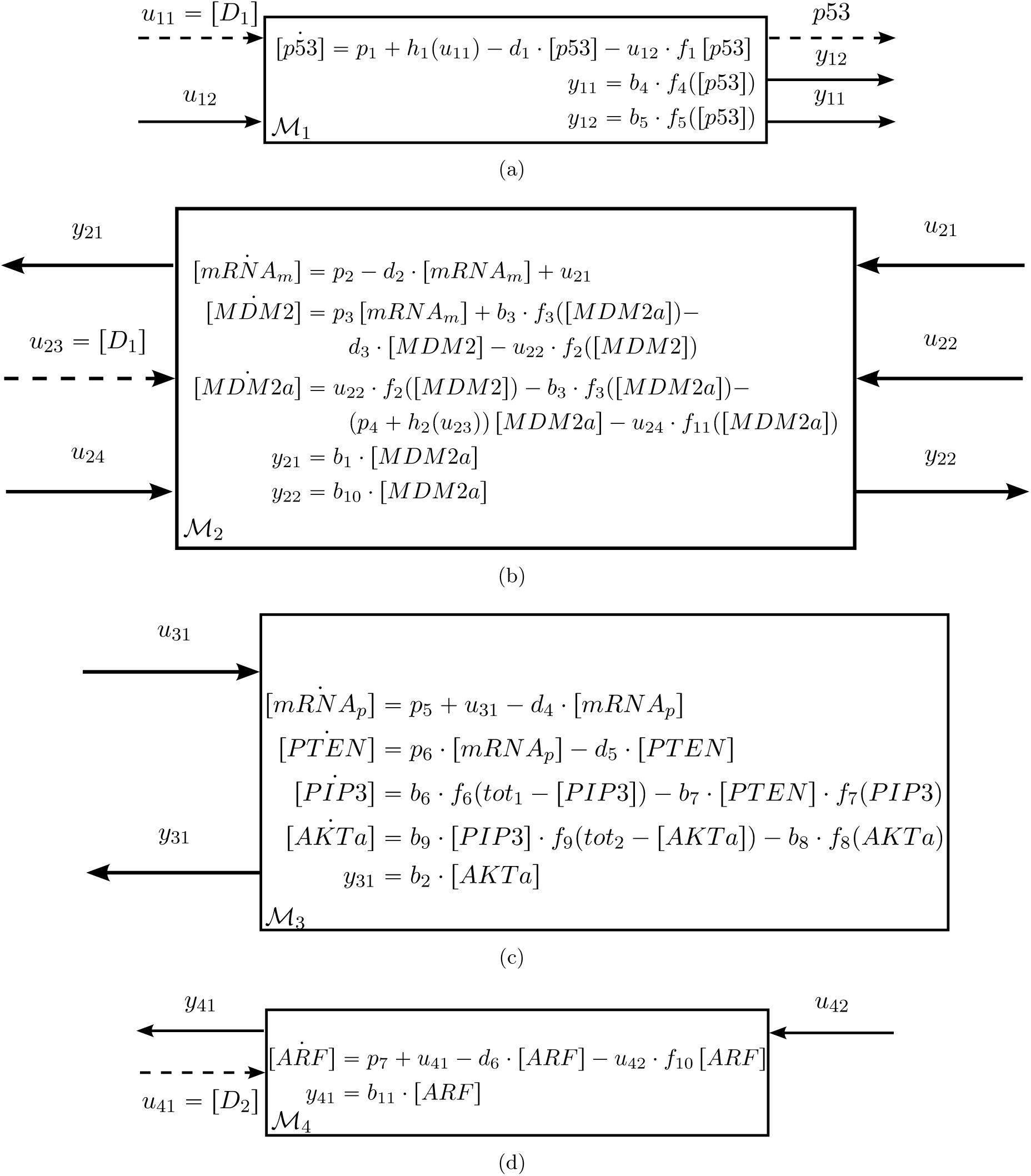
ODEs associated with each module (a) 𝒨_1_ (b) 𝒨_2_ (c) 𝒨_3_ (d) 𝒨_4_

Since there are multiple known *MDM*2 — *ARF* binding sites [15], we assume that these two interactions occur at separate binding sites, which results in the degradation rates of *MDM2a* given by *b*_11_ [*ARF*] *fn* [*MDM2a*], and that of *ARF* given by *b*_10_ [*MDM2*a] *f*_10_ [*ARF*], taking on distinct values.

In Fig. 2 of the main text, we showed a simplified block diagram representation of the full network 𝒩_υ_ consisting of the interconnections of the modules shown in S1. In Fig. S2, we show a more detailed block diagram representation of 𝒩_υ_. In addition, Fig. S3 shows the block diagram representations of the remaining different network configurations that we study in the main text.

**Figure S2:**
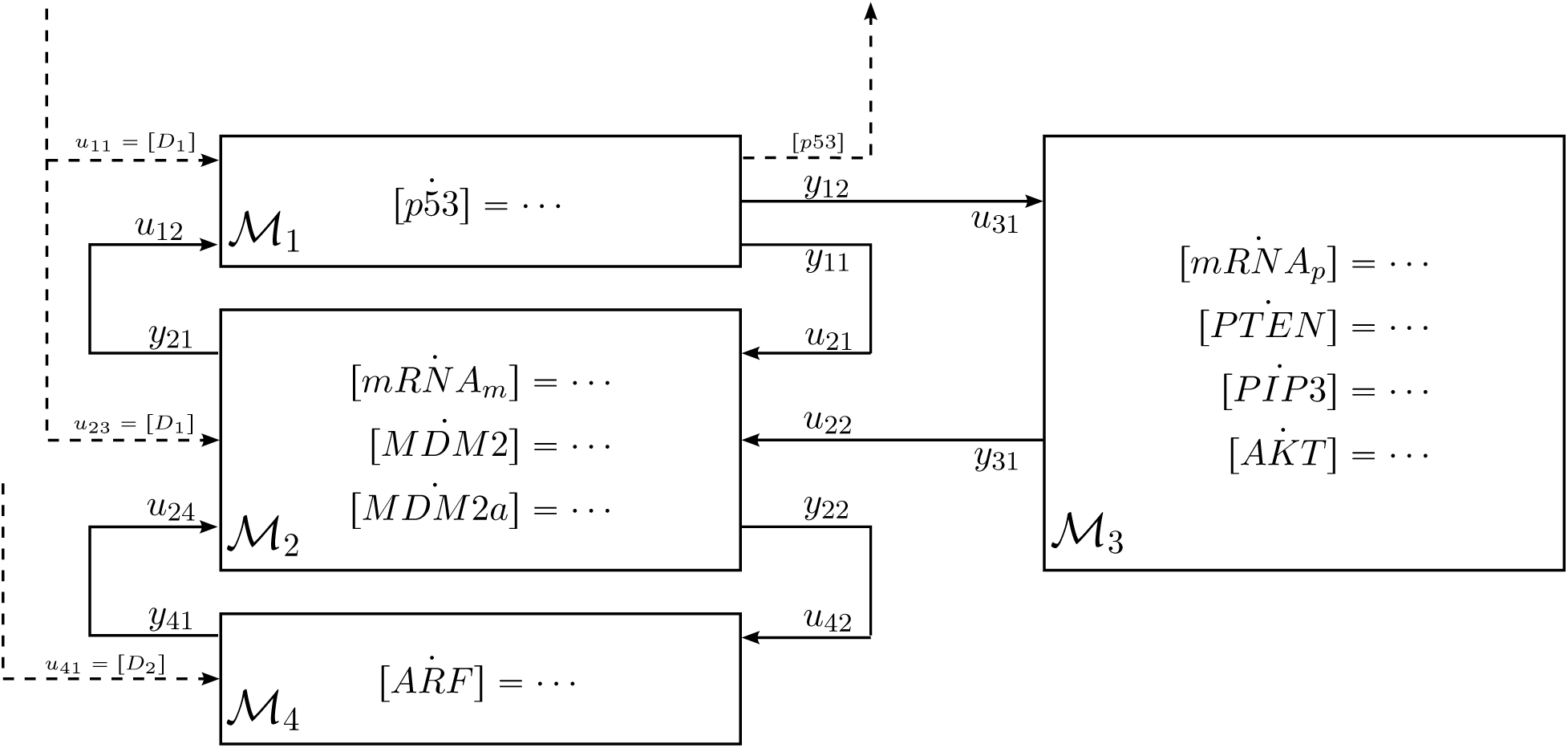
Modular block-diagram representations of full network 𝒩_υ_.

### A.2 Model parameters

The functions *f_i_*(*x*), *i* ∊ {1,2, …, 10,11} that appear in the models in Figure S1 are given by

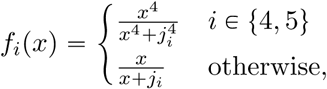

and the functions *h_i_*(*x*), *i* ∊ {1,2}, which represent the kinase activities, are given by

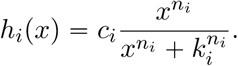

#### A.2.1 Tristable network 𝒩_*h*_

The parameters for this network are given in Table S1. Most parameters are consistent with the values and ranges from [6], except for the parameters relating to the *ARF-MDM*2 interactions and *ARF* reactions, that did not appear in [6]. The parameters relating to the entry of damage into the core regulation network through the kinases (*p*_1_, *p*_4_, *c*_1_, *c*_2_, *k*_1_ and *k*_2_) are chosen somewhat arbitrarily as was done in [6]. Moreover, while [6] chose to use *μM* as their unit of measurement, we used arbitrary unites (A.U.) in line with what is done in experimental studies [16, 17]. Since the amplitude of responses are known to vary significantly between different cells [18], the ratios between the parameters play a far more crucial role in bringing about the dynamical behavior as opposed to the parameters themselves. The steady-state solution to the network with these parameters results in the bifurcation diagram 6(a), which has been normalized with respect to the *p*53 level at *T*_0_.

#### A.2.2 Bistable networks 𝒩_2_ and 𝒩_3_

The study of 𝒩_2_ was performed with the same parameters as shown in Table S1, with the external input *u*_24_ set to 0 (although the results could be generalized to any value of *u*_24_). These parameters lead to the bifurcation diagram given by Fig. 4(a), which has been normalized by the *p*53 level at *T*_0_. To obtain the irreversible bistable switch, only five parameters corresponding to the entry of damage into the network were altered, and these are shown in Table S2.

**Table S1:** Parameters for the core regulation network model in the Network 𝒩_*h*_

| Parameter | Value | Units | Parameter | Value | Units |
| --- | --- | --- | --- | --- | --- |
| $b_1$ | 0.0625 | \h | $d_1$ | 0.02 | /h |
| $b_2$ | 22 | /h | $d_2$ | 0.01 | /h |
| $b_3$ | 0.5 | A.U./h | $d_3$ | 0.005 | /h |
| $b_4$ | 0.0375 | A.U./h | $d_4$ | 0.01 | /h |
| $b_5$ | 0.006 | A.U./h | $d_5$ | 0.0054 | /h |
| $b_6$ | 0.15 | A.U./h | $d_6$ | 0.001 | /h |
| $b_7$ | 73 | /h | $p_1$ | 0.017 | A.U./h |
| $b_8$ | 0.2 | A.U./h | $p_2$ | 0.0009 | A.U./h |
| $b_9$ | 20 | /h | $p_3$ | 0.02 | /h |
| $b_{10}$ | 1 | /h | $p_4$ | 0.0039 | /h |
| $b_{11}$ | 0.01 | /h | $p_5$ | 0.0009 | A.U./h |
| $j_1$ | 0.01 | A.U. | $p_6$ | 0.02 | /h |
| $j_2$ | 0.6 | A.U. | $p_7$ | 0.021 | A.U./h |
| $j_3$ | 0.1 | A.U. | $tot_1$ | 1 | A.U. |
| $j_4$ | 0.84 | A.U. | $tot_2$ | 1 | A.U. |
| $j_5$ | 1.19 | A.U. | $c_1$ | 0.066 | A.U./h |
| $j_6$ | 0.1 | A.U. | $c_2$ | 0.183 | A.U./h |
| $j_7$ | 0.5 | A.U. | $k_1$ | 2 | A.U. |
| $j_8$ | 0.1 | A.U. | $k_2$ | 3 | A.U. |
| $j_9$ | 0.1 | A.U. | $n_1$ | 1 | - |
| $j_{10}$ | 0.01 | A.U. | $n_2$ | 1 | - |
| $j_{11}$ | 0.01 | A.U. | | | |

**Table S2:** Parameters altered to obtain the irreversible bistable switch in the Network 𝒩_2_.

| Parameter | Value | Units |
| --- | --- | --- |
| $p_1$ | 0.0229 | A.U./h |
| $p_4$ | 0.0147 | /h |
| $c_1$ | 0.0559 | A.U./h |
| $c_2$ | 0.1191 | A.U./h |
| $k_2$ | 2 | A.U. |

The study of 𝒩_3_ was performed by slightly altering some parameters with respect to Table S1. These parameters are shown in Table S3. Most of the parameters altered are those which affect the entry of damage into the core regulation network through the kinases (*p*_1_, *p*_4_, *c*_1_ and *c*_2_), which were chosen arbitrarily. The strength of the negative feedback between *MDM*2 and *p*53 is strengthened by increasing the parameter *b*_1_. This does not qualitatively change the network behavior other than to clearly demarcate the higher and lower branches of *p*53 equilibrium points by increasing the distance between these branches. The external input *u*_22_ to this network is set to 0.1.

The bifurcation diagram of this network is shown in Fig. S4. This diagram was drawn assuming that [*D*_1_] and [*D*_2_] are coupled, which is not unreasonable since they are both triggered by DNA double-strand breaks.

**Table S3:** Parameters altered for the core regulation network model in the Network 𝒩_3_

| Parameter | Value | Units |
| --- | --- | --- |
| $b_1$ | 1.25 | $\backslash h$ |
| $p_1$ | 0.017 | A.U./h |
| $p_4$ | 0.0039 | /h |
| $c_1$ | 0.066 | A.U./h |
| $c_2$ | 0.183 | A.U./h |

**Figure S4:**
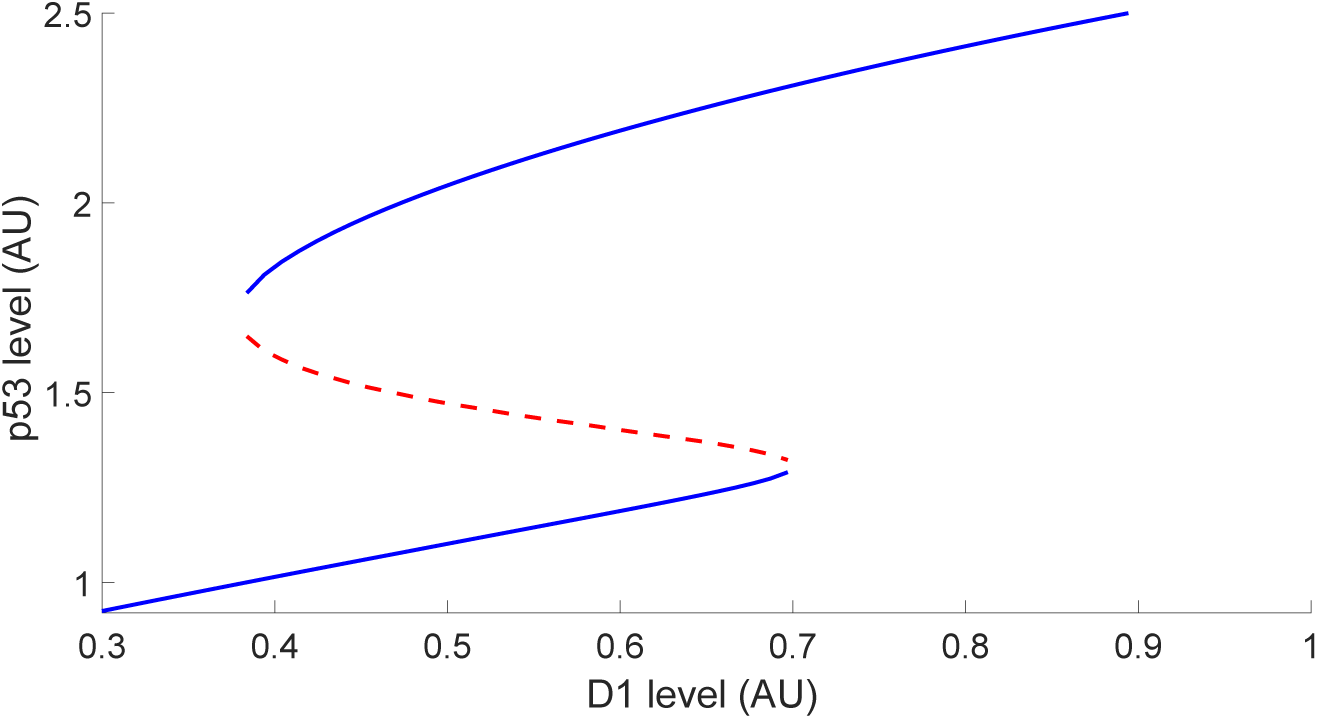
Bifurcation diagram for the network 𝒩_3_.

In the main text, we showed simulations of the full network in response to varying levels of damage. We stated that the reversible bistable network is able to admit both pulsing for moderate damage and a monotonic switch in *p*53 to a high level for high damage, but that this switch was reversible. We demonstrated this through a simulation of the network 𝒩_2_ with an initial *#DSB =* 450, and illustrate our result in Fig. S5.

**Figure S5:**
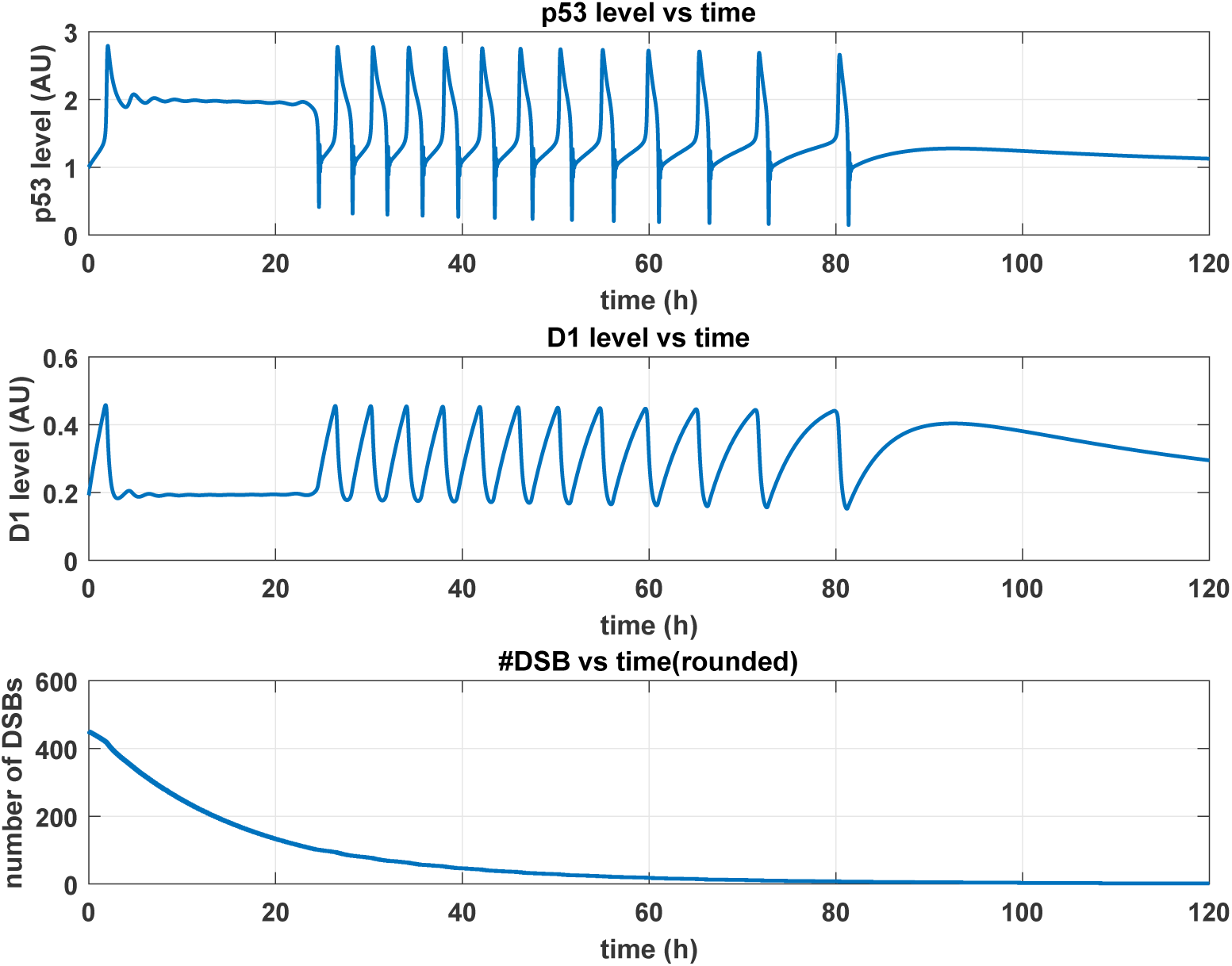
*p*53 and *D*_1_ responses, along with DSB repair process for initial *#DSB* = 450 in 𝒩_2_.

### A.3 Monostable networks 𝒩_1_ and 𝒩_*i*_

The study of the networks 𝒩_1_ and 𝒩_*i*_ was performed with the parameters shown in Table S1, except for those that affect the effect of DNA damage in the core regulation network through the kinases (*p*_1_, *p*_4_, *c*_1_ and *c*_2_). The parameters were chosen to illustrate how these networks could operate as ultrasensitive switches, and are given in Table S4. The resulting *p*53 responses to changes in [*D*_1_] are seen in Fig. 3, with the y-axis normalized by the initial concentration of *p*53.

**Table S4:** Parameters altered for the core regulation network models in Networks 𝒩_i_ and 𝒩_i_.

| Parameter | Value | Units |
| --- | --- | --- |
| $p_1$ | 0.0206 | A.U./h |
| $p_4$ | 0.1 | /h |
| $c_1$ | 0.0226 | A.U./h |
| $c_2$ | 0.0531 | A.U./h |
| $k_1$ | $2.79 * 10^{-7}$ | A.U. |
| $k_2$ | $2.79 * 10^{-7}$ | A.U. |
| $n_1$ | 17 | - |
| $n_2$ | 17 | - |

## B Damage sensing and repair model

The functions for Eqs. (1)–(3) of the main text are given by

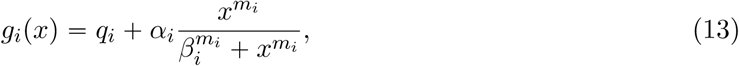

with the parameters given in Table S5.

**Table S5:**
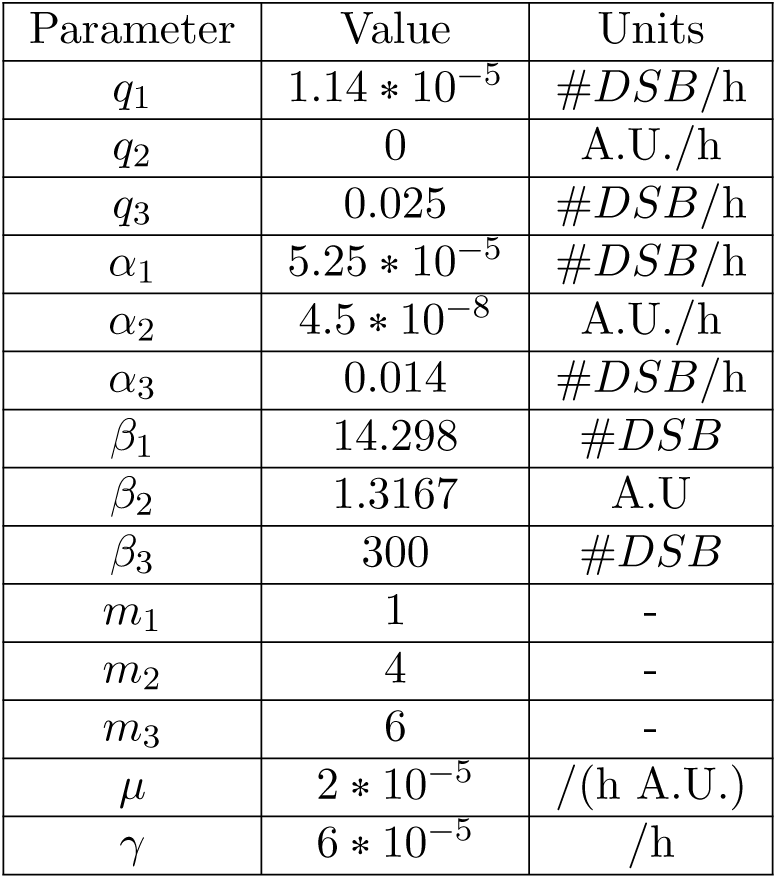
Parameters altered for core regulation network models Network 𝒩_1_ and 𝒩_i_.

As was noted in the main text, the dynamics of *D*_1_ are assumed to be slow compared to that of the core regulation network, leading to the observed behavior. We simplified our simulations by linearizing *g*_2_ in the following way:

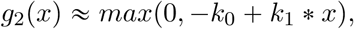

where

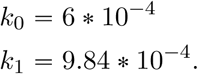

## C Algorithm to compute network equilibrium points

In this section, we describe the algorithm used to compute the equilibrium points of our networks and obtain the bifurcation diagrams.

### C.1 The Steady-State Behavior of a module

The procedure that the algorithm employs involves *gridding* the equilibrium values of the input and output signals for each module to form a table known as the *Steady-State Behavior* (SSB) of a module [19]. Typically, this table is obtained by computing the equilibrium values of the outputs for a range of constant inputs, and this special case of the SSB has been termed the *Static Characteristic Function* (SCF) [20, 21] of the module. It is worth noting that these tables could be multivalued, as there could be multiple output equilibria for a given input as is evidenced in the bistable and tri-stable networks.

The number of degrees of freedom of each module is the total number of inputs to that module. Given a module with *p* inputs and *q* outputs, instead of gridding over the range of *p* inputs, it is often more convenient to grid the *p* degrees of freedom by gridding over a combination of inputs and outputs. We illustrate this using 𝒨_1_, where the equilibrium equation is given by

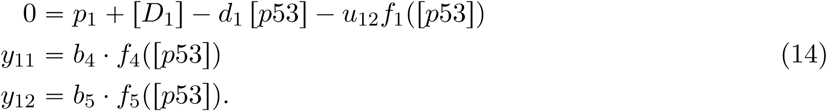

Given that *f*_1_ is a Hill function of [*p*53] (see Section A.2), it is more straightforward to solve for *u*_12_ as a function of [*p*53] and [*D*_1_]. We can then obtain a table of equilibrium values of the signals, that can be formally represented by the SSB of the module as

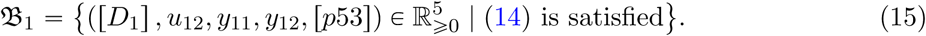

A tabular representation of (15) is illustrated in Table S6.

**Table S6:** Table of equilibrium values for 𝒨_1_. The superscript *i* denotes the *i^th^* row of the table.

| $[D_1]$ | $[p53]$ | $u_{12}$ | $y_{11}$ | $y_{12}$ |
| --- | --- | --- | --- | --- |
| ⋮ | ⋮ | ⋮ | ⋮ | ⋮ |
| $[D_1]^i$ | $[p53]^i$ | $u_{12}^i$ | $y_{11}^i$ | $y_{12}^i$ |
| ⋮ | ⋮ | ⋮ | ⋮ | ⋮ |

We can obtain the SSBs of the other modules as well, denoted by 𝔅_2_ − 𝔅_4_ for modules 𝒨_2_ – 𝒨_4_ respectively, with the corresponding tabular representations shown in Tables S7-S9.

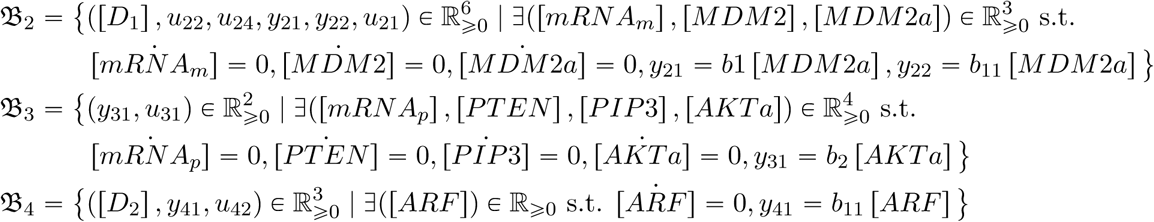

**Table S7:** Table of equilibrium values for 𝒨_2_. The superscript *j* denotes the *j^th^* row of the table.

| $[D_1]$ | $u_{22}$ | $u_{24}$ | $y_{21}$ | $y_{22}$ | $u_{21}$ |
| --- | --- | --- | --- | --- | --- |
| ⋮ | ⋮ | ⋮ | ⋮ | ⋮ | ⋮ |
| $[D_1]^j$ | $u_{22}^j$ | $u_{24}^j$ | $y_{21}^j$ | $y_{22}^j$ | $u_{21}^j$ |
| ⋮ | ⋮ | ⋮ | ⋮ | ⋮ | ⋮ |

**Table S8:** Table of equilibrium values for 𝒨_3_. The superscript *k* denotes the *k^th^* row of the table.

| $y_{31}$ | $u_{31}$ |
| --- | --- |
| ⋮ | ⋮ |
| $y_{31}^k$ | $u_{31}^k$ |
| ⋮ | ⋮ |

**Table S9:**
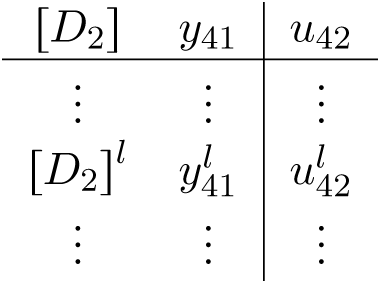
Table of equilibrium values for 𝒨_4_. The superscript l denotes the *k^th^* row of the table.

| $[D_2]$ | $y_{41}$ | $u_{42}$ |
| --- | --- | --- |
| ⋮ | ⋮ | ⋮ |
| $[D_2]^l$ | $y_{41}^l$ | $u_{42}^l$ |
| ⋮ | ⋮ | ⋮ |

### C.2 A modular approach to computing the equilibrium point of a network

To solve for the equilibrium value of [*p*53] of the network 𝒩_υ_ as a function of [*D*_1_] and [*D*_2_], we need to systematically eliminate the input and output signals flowing between the modules, also known as the *latent variables* in the network [19]. We demonstrate this by focusing on the interconnection between Modules *M*_1_ and *M*_2_ as is shown in Fig. S2, where we see the associations *y*_11_ = *u*_21_ and *y*_21_ = *u*_12_. To eliminate these latent variables, we identify the rows *i* and *j* in Tables S6 and S7 respectively such that

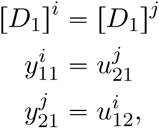

leading to the Table S10. Mathematically, the construction of Table S10 corresponds to computing the SSB set 𝔅_12_, where

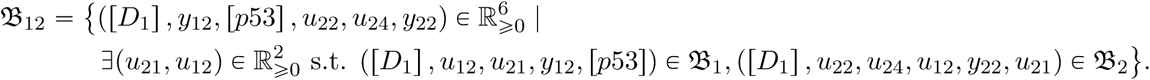

Computationally, the construction of Table S10 is achieved by finding the intersection points between each of the lines mapping from *u*_12_ to *y*_11_ from Table S6, and each of the lines mapping from *u*_21_ to *y*_21_ from Table S7, for all combination of other variables, as outlined in Fig. S6.

**Table S10:**
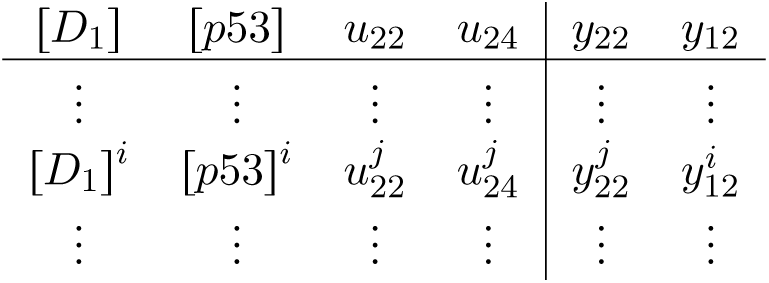
Table of equilibrium values when modules Mi and M2 are combined.

| $[D_1]$ | $[p53]$ | $u_{22}$ | $u_{24}$ | $y_{22}$ | $y_{12}$ |
| --- | --- | --- | --- | --- | --- |
| ⋮ | ⋮ | ⋮ | ⋮ | ⋮ | ⋮ |
| $[D_1]^i$ | $[p53]^i$ | $u_{22}^j$ | $u_{24}^j$ | $y_{22}^j$ | $y_{12}^i$ |
| ⋮ | ⋮ | ⋮ | ⋮ | ⋮ | ⋮ |

**Figure S6:**
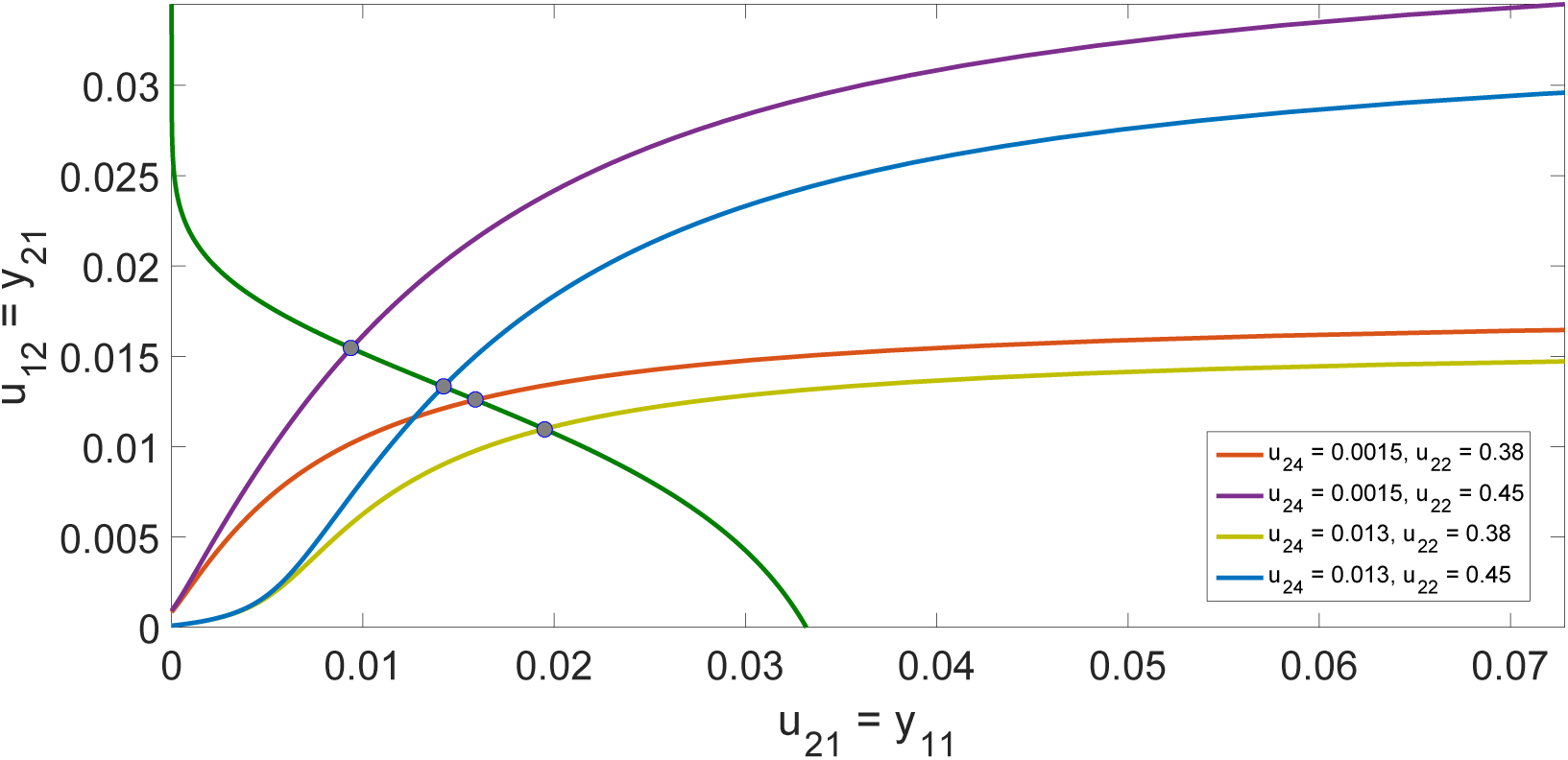
Projections of Table S6 and Table S7 into the *u*_12_ vs *y*_11_ (green line) and *y*_21_ vs *u*_21_ (other lines) planes respectively, for different values of u_24_ and *u*_22_ with [*D*_1_] = 0.4. The gray intersection points correspond to the equilibrium values of *u*_12_ and *u*_21_ for the given *u*_24_, *u*_22_ and [*D*_1_].

The next step is to identify the rows *m* and *k* from Tables S10 and S8 respectively such that

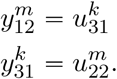

The elimination of these latent variables results in Table S11, whose construction can be mathematically represented by the SSB

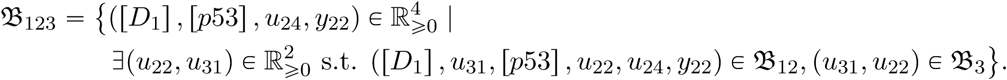

Computationally, the construction of Table S11 is achieved by finding the intersection points between each of the lines mapping from *u*_22_ to *y*_12_ from Table S10, and each of the lines mapping from *y*_31_ to *u*_31_ from Table S8, for all combination of other variables, as outlined in Fig. S7.

**Figure S7:**
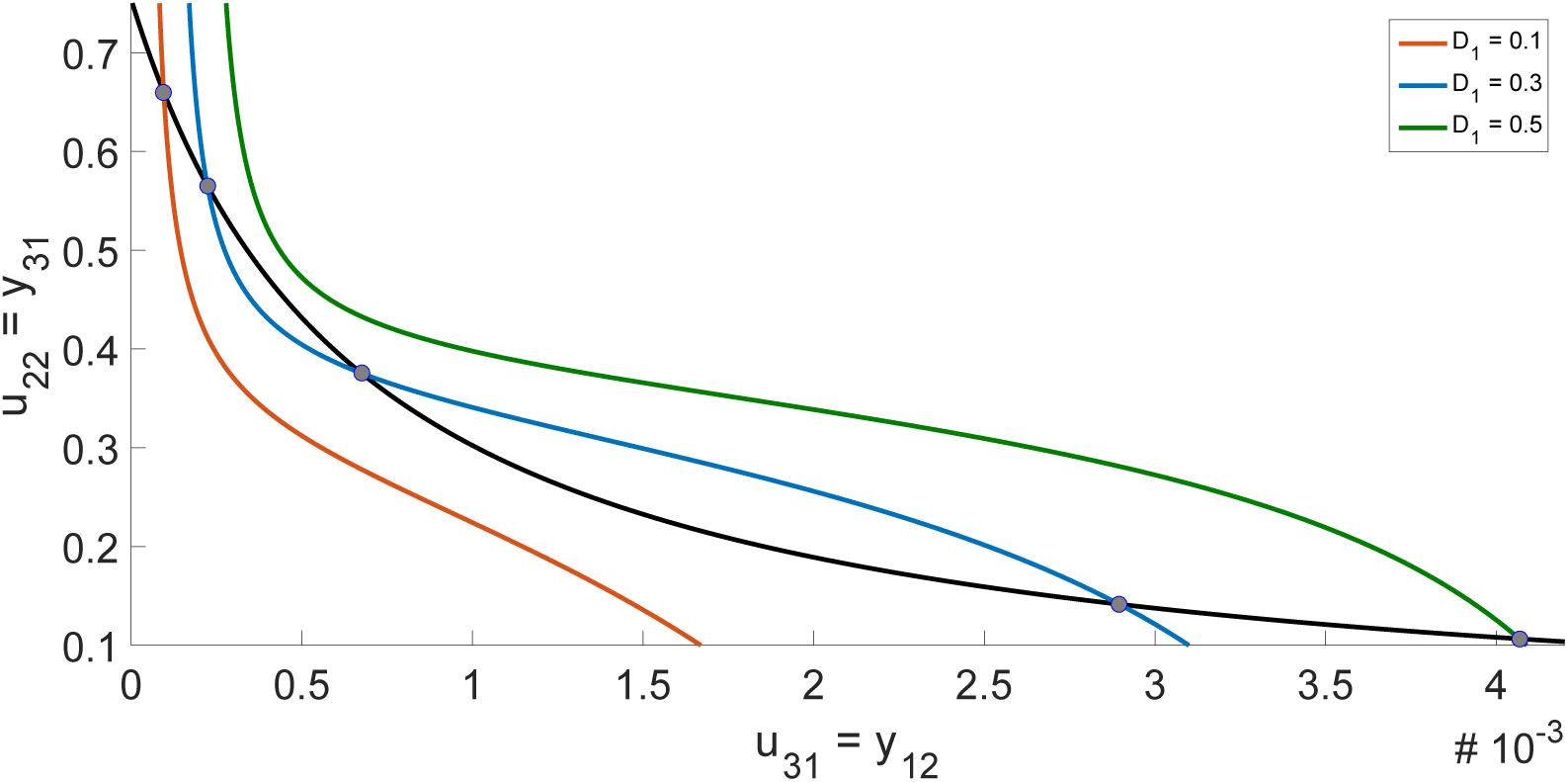
Projections of Table S10 and Table S8 into the *u*_22_ vs *y*_12_ (black line) and *y*_31_ vs *u*_31_ (other lines) planes respectively, for different values of [*D*_1_] with *u*_24_ = 0.0015. The gray intersection points correspond to the equilibrium values of *u*_31_ and *u*_22_ for the given *u*_24_ and [*D*_1_]. The bifurcation can clearly be seen as [*D*_1_] increases, with [*D*_1_] = 0.1 and [*D*_1_] = 0.5 corresponding to a single equilibrium point, while [*D*_1_] = 0.3 corresponds to three equilibrium points.

**Table S11:** Table of equilibrium values when modules 𝒨_1_, 𝒨_2_ and 𝒨_3_ are combined.

| $[D_1]$ | $[p53]$ | $u_{24}$ | $y_{22}$ |
| --- | --- | --- | --- |
| ⋮ | ⋮ | ⋮ | ⋮ |
| $[D_1]^m$ | $[p53]^m$ | $u_{24}^m$ | $y_{12}^m$ |
| ⋮ | ⋮ | ⋮ | ⋮ |

Finally, we identify the rows *n* and *l* from Tables S11 and S9 respectively such that

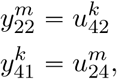

The elimination of these latent variables results in Table S12, which is the combined steady-state behavior of the entire network 𝒩_υ_. Mathematically, Table S12 corresponds to the set

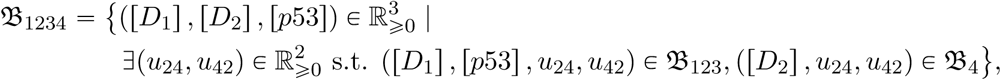

which was precisely what we wanted to compute. Computationally, the construction of Table S11 is achieved by finding the intersection points between each of the lines mapping from *u*_24_ to *y*_22_ from Table S11, and each of the lines mapping from *y*_41_ to *u*_41_ from Table S8, for all combination of [*D*_1_] and *[*D*_2_]*, as outlined in Fig. S8.

**Figure S8:**
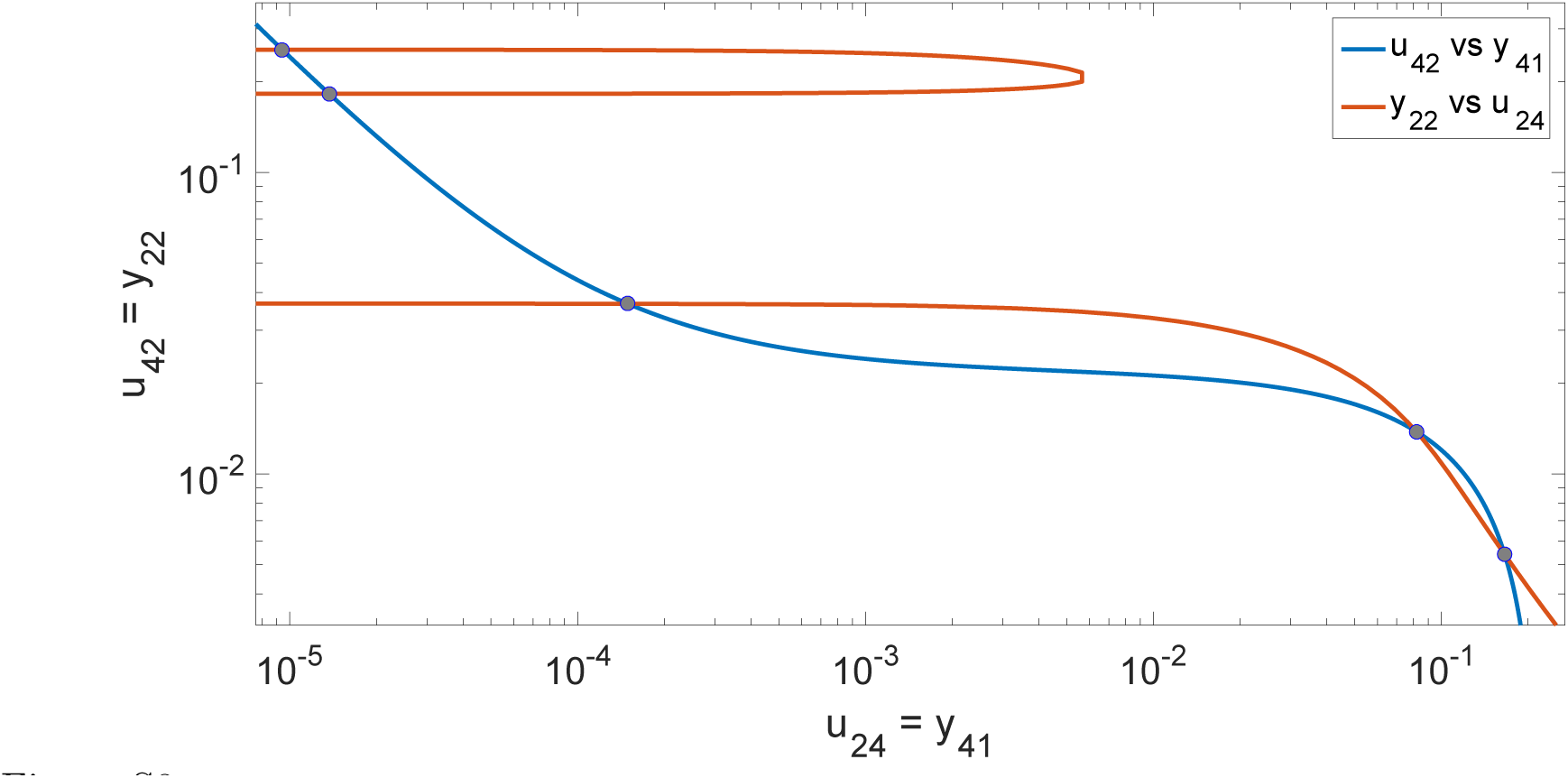
Projections of Table S11 and Table S9 into the *y*_22_ vs *u*_24_ (red line) and *u*_42_ vs *y*_41_ (blue line) planes respectively, for different values of [*D*_1_] = 0.3 with [*D*_2_] = 0.002. The gray intersection points correspond to the equilibrium values of *u*_42_ and *u*_24_ for the given [*D*_1_] and [*D*_2_]. The equilibrium values can be found for a range of [*D*_1_] and [*D*_2_], and then used to compute the corresponding *p*53 equilibrium values.

**Table S12:** Table of equilibrium values for the entire network 𝒩_υ_.

| $[D_1]$ | $[D_2]$ | $[p53]$ |
| --- | --- | --- |
| ⋮ | ⋮ | ⋮ |
| $[D_1]^n$ | $[D_2]^l$ | $[p53]^n$ |
| ⋮ | ⋮ | ⋮ |

The process of computing intersection points between lines was done using a MATLAB script available on-line [22]. The algorithm described above can be generalized for arbitrary networks, and work is ongoing to identify the complexity of this algorithm for large networks.

The algorithm described above allows us to compute the equilibrium points. The stability of each point is then determined through an eigenvalue analysis of the network linearized about each equilibrium point.

**Figure S3:**
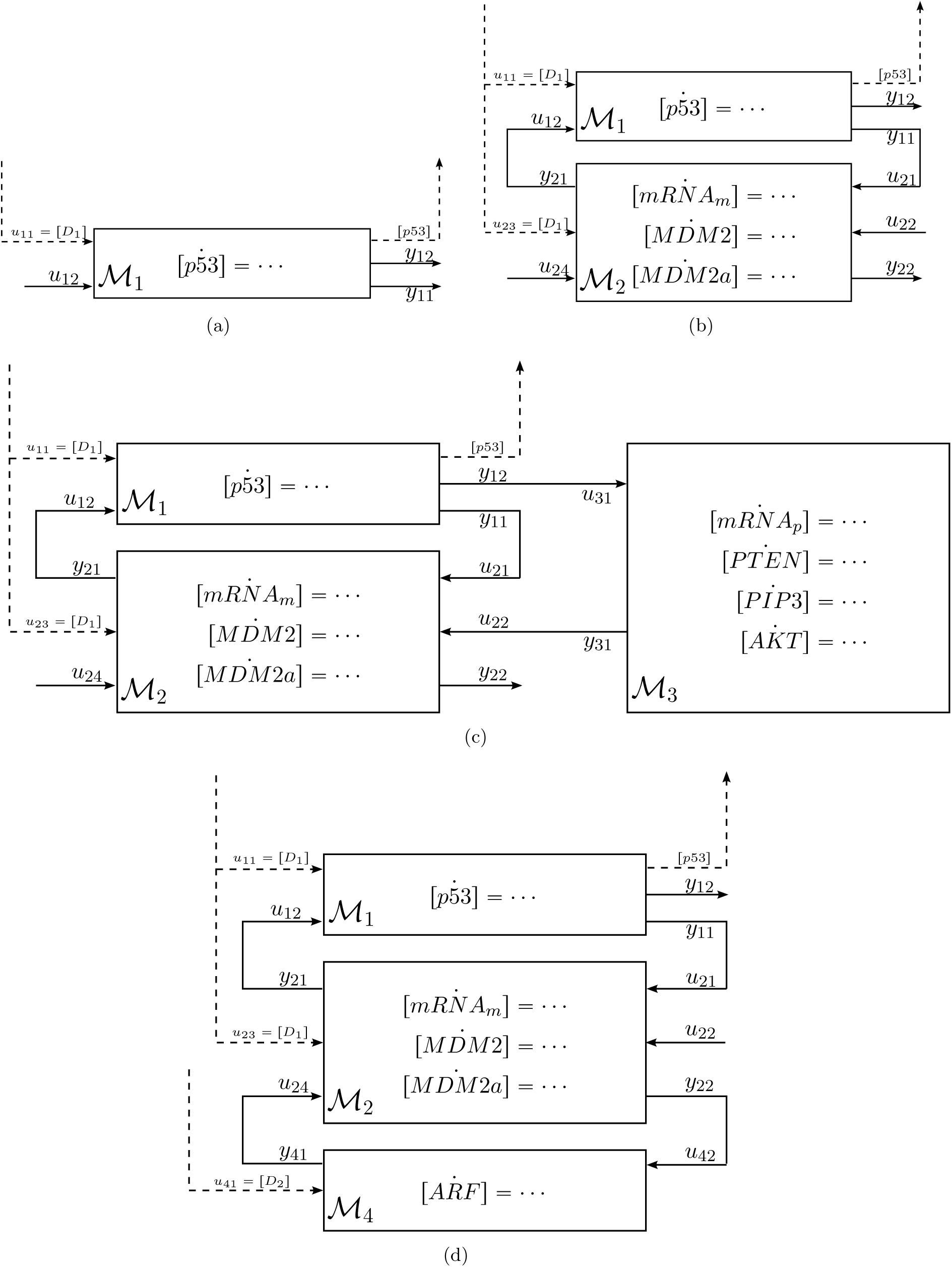
Modular block-diagram representations of the various *p*53 pathway configurations found in the evolutionary data. The specific ODEs describing each module are shown in Fig. S1. (a) Network 𝒩_*i*_ (b) Network 𝒩_1_ (c) Network 𝒩_2_ (d) Network 𝒩_3_

